# Immobilised collagen prevents shedding and induces sustained GPVI clustering and signalling in platelets

**DOI:** 10.1101/2020.07.07.192435

**Authors:** Chiara Pallini, Jeremy A. Pike, Christopher O’Shea, Robert K. Andrews, Elizabeth E. Gardiner, Steve P. Watson, Natalie S. Poulter

## Abstract

Collagen, the most thrombogenic constituent of blood vessel walls, activates platelets through glycoprotein VI (GPVI). In suspension, following platelet activation by collagen, GPVI is cleaved by A Disintegrin And Metalloproteinase (ADAM)10 and ADAM17. In this study, we use single-molecule localization microscopy and a 2-level DBSCAN-based clustering tool to show that GPVI remains clustered along immobilised collagen fibres for at least 3 hours in the absence of significant shedding. Tyrosine phosphorylation of spleen tyrosine kinase (Syk) and Linker of Activated T cells (LAT), and elevation of intracellular Ca^2+^, are sustained over this period. Syk, but not Src kinase-dependent signalling is required to maintain clustering of the collagen integrin α2β1, whilst neither is required for GPVI. We propose that clustering of GPVI on immobilised collagen protects GPVI from shedding in order to maintain sustained Src and Syk-kinases dependent signalling, activation of integrin α2β1 and continued adhesion.

## Introduction

Platelets are small anucleate blood cells which play important roles in many physiological and pathological processes including haemostasis, thrombosis, inflammation and cancer(*1*). Collagen, one of the major components of the extracellular matrix (ECM), is a potent activator of platelets acting through its interaction with the immunoglobulin receptor glycoprotein VI (GPVI). GPVI is considered to be a promising anti-thrombotic drug target due to its restricted expression pattern in platelets and megakaryocytes and relatively minor role in haemostasis(*2*). More recently GPVI has been shown to be involved in maintaining vascular integrity at sites of inflammation(*3*).

GPVI is an ∼65 kDa transmembrane protein found at the platelet surface in association with the Fc receptor gamma chain (FcRγ). Ligand engagement of GPVI triggers Src family kinase-dependent phosphorylation of the conserved tyrosines in the immunoreceptor tyrosine-based activation motif (ITAM) in the associated FcRγ chain and binding of the spleen tyrosine kinase (Syk) via its tandem SH2 domains(*4-6*). Activation of Syk, by Src-dependent phosphorylation and autophosphorylation, initiates formation of a Linker of Activated T cells (LAT)-based signalosome that culminates in the activation of phospholipase Cγ2 (PLCγ2), elevation of intracellular Ca^2+^, integrin activation (including the collagen-binding integrin α2β1) and platelet activation(*7*). The GPVI-FcRγ complex is a signalling rather than an adhesive receptor complex with activation of integrins required for stable platelet adhesion under flow(*8*).

It has recently been demonstrated that GPVI is also a receptor for fibrin(*9-12*) and fibrinogen(*10, 13*), with ligand binding also resulting in platelet activation. Different ligands induce different degrees of higher order clustering of GPVI thereby regulating signal strength(*14-17*). In suspension, activation of platelets by collagen and other ligands leads to shedding of GPVI through the action of A Disintegrin And Metalloproteinase (ADAM)10 and ADAM17(*18*), thereby limiting activation. Paradoxically, this could potentially lead in vivo to the unwanted embolisation, or breaking away, of single platelets and platelet aggregates which could lead to further complications. To date, shedding of GPVI in platelets adhered to and spreading on an immobilised collagen substrate, which would represent the first steps of thrombus formation or adhesion of platelets to the ECM in a blood vessel with compromised integrity, has not been investigated.

In the present study, we have used single-molecule localization microscopy (SMLM) to monitor clustering of GPVI on platelets adhering and spreading on immobilised fibrillar collagen and correlated this with biochemical data and other complementary imaging techniques to assess platelet activation and GPVI shedding. We show that, in contrast to studies in solution, GPVI is not shed from platelets activated by immobilised collagen and that signalling is sustained, with Syk, LAT and Ca^2+^ mobilisation remaining active over several hours of platelet spreading. GPVI clustering is not reversed by inhibition of Src or Syk kinase activation, in contrast to integrin α2β1, which requires sustained Syk but not Src-dependent signalling, highlighting the different nature of the two collagen receptors. We further show that the metalloproteinase ADAM10 does not co-localise with the GPVI clustered at collagen fibrils, suggesting exclusion of the sheddase from the clusters. Our findings confirm the hypothesis that sustained GPVI signalling is further regulated by higher-order clustering, acting to protect GPVI from shedding, essential for prolonged platelet activation, including integrin activation which is required to maintain platelet spreading. This is likely to be important for thrombus stability and the prevention of bleeding associated with disrupted vascular integrity driven by inflammation.

## Results

### GPVI signalling is sustained over time in platelets adhering to immobilised collagen

Studies in vitro using platelets and cell lines activated with GPVI receptor agonists in suspension have demonstrated that fibrillar collagen triggers slow but sustained ITAM signalling(*19*) and metalloproteolytic shedding of GPVI(*18*). This raises the question of whether this is also the case on an immobilised surface, where shedding might lead to ablation of adhesion to collagen and embolisation of platelets. Reflection images show that human platelets remain spread on immobilised collagen for at least 3 h (Fig. 1A). As expected, there was a steady increase in the number of platelets coming into contact with the collagen fibres over time, but there was no change in mean platelet surface area (Fig. 1B, C). To assess platelet signalling in these cells, whole cell tyrosine phosphorylation and specific phosphorylation sites on two key GPVI signalling mediators, Syk and LAT, were measured by western blot (Fig. 1D). Total phospho-tyrosine levels, detected by the antibody clone 4G10, were elevated in adherent platelets relative to cells in suspension (non-adhered, NA) and remained increased over time (Fig. 1D). Site-specific phosphorylation of both Syk Tyr ^525/6^ and LAT Tyr^200^, residues associated with their activation, was maintained, with no significant change in signal over 3 h (Fig. 1D and Supp. Fig. 1A, B). An additional step of GPVI signalling transduction involves Ca^2+^ movement from intracellular compartments upon activation of PLCγ2. Therefore, we monitored the Ca^2+^ spiking in platelets labelled with the Ca^2+^ dye Oregon green 488 BAPTA-1-AM that had been spread on immobilised collagen for up to 3 h (Supp. movie 1). Single cell calcium dynamics was analysed using MATLAB. The results show that the percentage and frequency of spiking platelets spread on collagen for 3 h remained constant (Fig. 1E, F), although the amplitude and peak duration slightly declined (Fig. 1G, H). Overall, immobilised collagen supports platelet spreading via GPVI signalling for at least 3 h.

**Fig. 1.**
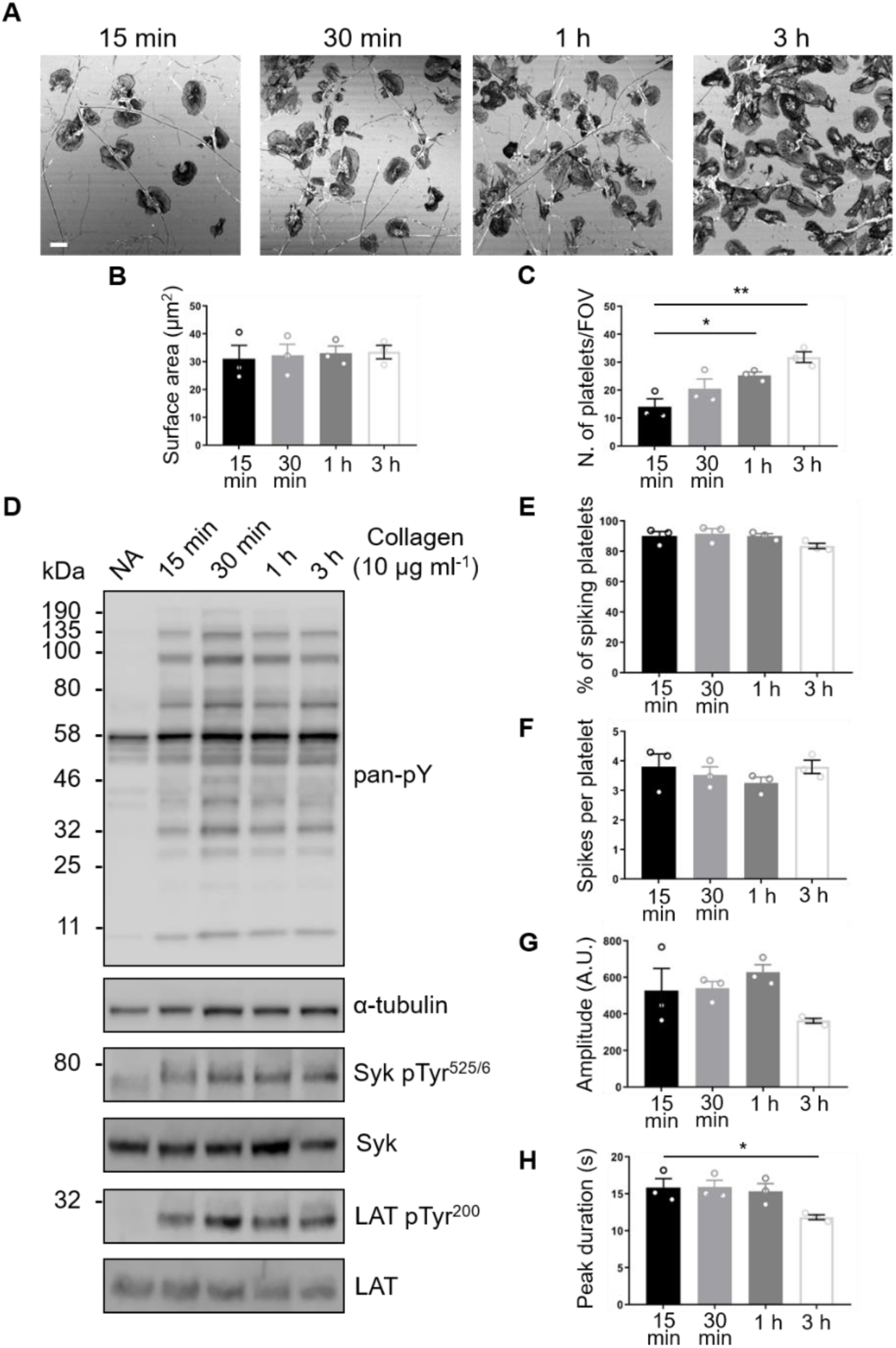
GPVI signalling remains constant in platelets spread on fibrous collagen for at least 3 h. (A) Confocal reflection images of washed human platelets spread on Horm collagen for the indicated times. Quantification of spread platelet surface area (B) and platelet number per field of view (C) for the specified time points from 5 FOVs from three independent experiments. Scale bar: 5 μm. (D) Western blot analysis of platelets spread for the indicated time points or non-adhered platelets (NA) probed for total phosphotyrosine (4G10), phosphorylated Syk (Tyr525/6; pSyk) and LAT (Tyr200; pLAT). The antibodies against α-tubulin, pan-Syk and pan-LAT were used as loading controls for total phosphotyrosines, phospho-Syk and phospho-LAT, respectively. Quantification of live-cell calcium imaging of platelets spread for the indicated time points showing percentage of platelets exhibiting calcium spiking (E), number of spikes per platelet in the 2 min imaging period (F), spike amplitude (G), spike duration (H). Data taken from three representative FOVs for each time point (∼150 platelets in each FOV) from three independent experiments. All data are expressed as mean ± SEM. Significance was calculated using one-way ANOVA with Tukey’s multiple comparisons test (* p < 0.05).

### Sustained signalling colocalises with GPVI on collagen in spread platelets

Western blot analysis quantifies platelet signalling events but does not provide spatial information. To investigate the location of signalling events, platelets spread on immobilised collagen for 1 h were labelled for either pan-Syk, phospho-Syk Tyr^525/6^, pan-LAT or phospho-LAT Tyr^200^ and imaged by confocal microscopy (Fig. 2A). Fig. 2Ai, iii illustrate that total (pan) Syk and LAT are homogenously distributed over the platelet surface. However, both pSyk and pLAT were enriched along collagen fibrils, with a lower density of phospho-proteins detected in areas where there is no visible collagen (Fig. 2Aii, iv). Dual-colour confocal imaging of GPVI and phosphorylated proteins was carried out to assess their colocalisation (Fig. 2B). GPVI, labelled with 1G5-Fab, was highly enriched along collagen fibres (Fig. 2Bi). A combination of qualitative (colocalisation mask, showing pixels which contain both GPVI and phospho-proteins, Fig. 2Biv) and quantitative (Pearson’s correlation coefficient, Fig. 2C) analysis demonstrated significant colocalisation between the phospho-proteins and GPVI along collagen. This colocalisation was sustained for at least 3 h, with the pLAT/GPVI colocalisation slighty, but significantly, increased at the later time point.

**Fig. 2.**
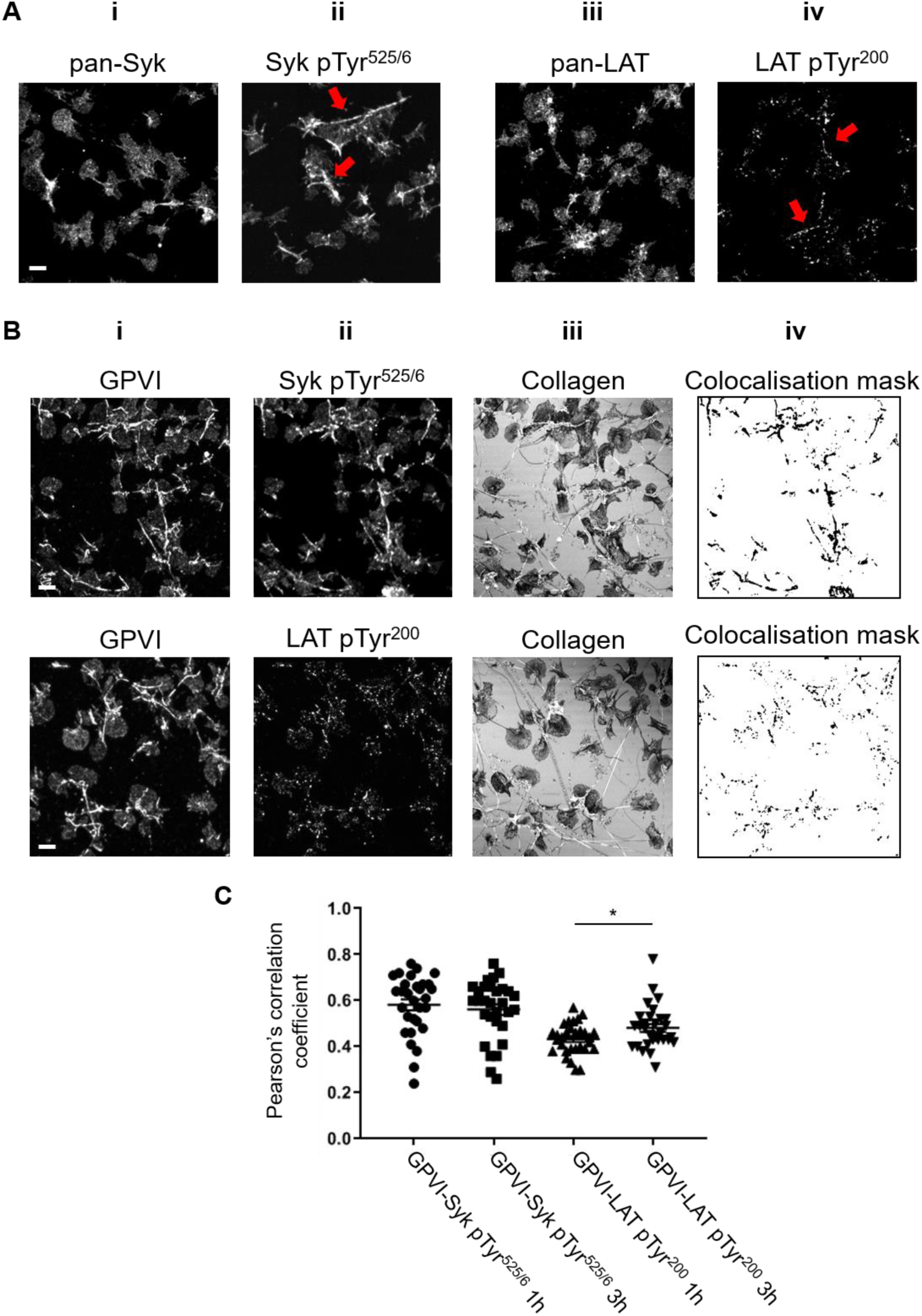
Phosphorylated Syk and LAT colocalise with GPVI along collagen fibres. Confocal microscopy imaging of human washed platelets spread on collagen (10 µg ml^-1^) for 1 h, labelled for (**Ai**) Pan-Syk, (**Aii**) pSyk Tyr^525/6^, (**Aiii**) Pan-LAT, (**Aiv**) pLAT Tyr^200^. The enrichment of pSyk Tyr^525/6^ and pLAT Tyr^200^ along collagen fibres is indicated by red arrows (**Aii, iv**). Dual-colour confocal imaging of platelets, pre-incubated for Pan-GPVI using 1G5-Fab (**Bi**), spread on collagen for 1 h and post-labelled for pSyk Tyr^525/6^ (**Bii**, first panel) or pLAT Tyr^200^ (**Bii**, second panel). The position of collagen fibres is shown in the confocal reflection image (**Biii**). The qualitative colocalisation mask in (**Biv**) shows the colocalisation of GPVI and the respective phosphoprotein with enrichment at the collagen fibres. (**C**) Quantification of GPVI and phosphoprotein colocalisation using Pearson’s correlation coefficient for platelets spreading for 1 h and 3 h. Scatter plot represents mean ± SEM of n=30 platelets taken from three independent experiments. Significance was measured using unpaired two-tailed t-test, * p < 0.05. Scale bar: 5μm.

### GPVI clustering and nanoclustering on collagen fibres is sustained over time

In order to correlate GPVI clustering with the platelet signalling events detected using biochemical methods, we used the SMLM technique dSTORM (characterized by a resolution of 20-40 nm)(*20*). dSTORM permitted the nanoscale spatial organisation of GPVI receptors in platelets that have been spread on collagen fibres for 1 h and 3 h to be examined and quantified relatively between conditions. The diffraction-limited TIRF images (Fig. 3Ai) show the same alignment of GPVI receptors along collagen fibres as that observed in confocal imaging (Fig. 2Bi). dSTORM allowed us to super-resolve the receptor distribution on fibrous collagen (Fig. 3Aii). A customised two-level cluster analysis workflow based on DBSCAN(*21*) was implemented to first identify and isolate large clusters of GPVI along collagen fibres (Level I; Fig. 3Aiii) and then take those clusters and subdivide them into nanoclusters (Level II; Fig. 3Aiv). This approach allowed us to assess only GPVI clusters associated with the visible collagen fibres, to identify any differences between the platelets spread for different durations. The algorithm clusters the localised data points by grouping detections with high local density. Different clusters are represented by different colours (Fig. 3Aiii, iv). Quantitative analysis of the clusters showed no significant difference in cluster area between 1 h and 3 h of spreading at either Level I or Level II clustering (Fig. 3B).

**Fig. 3.**
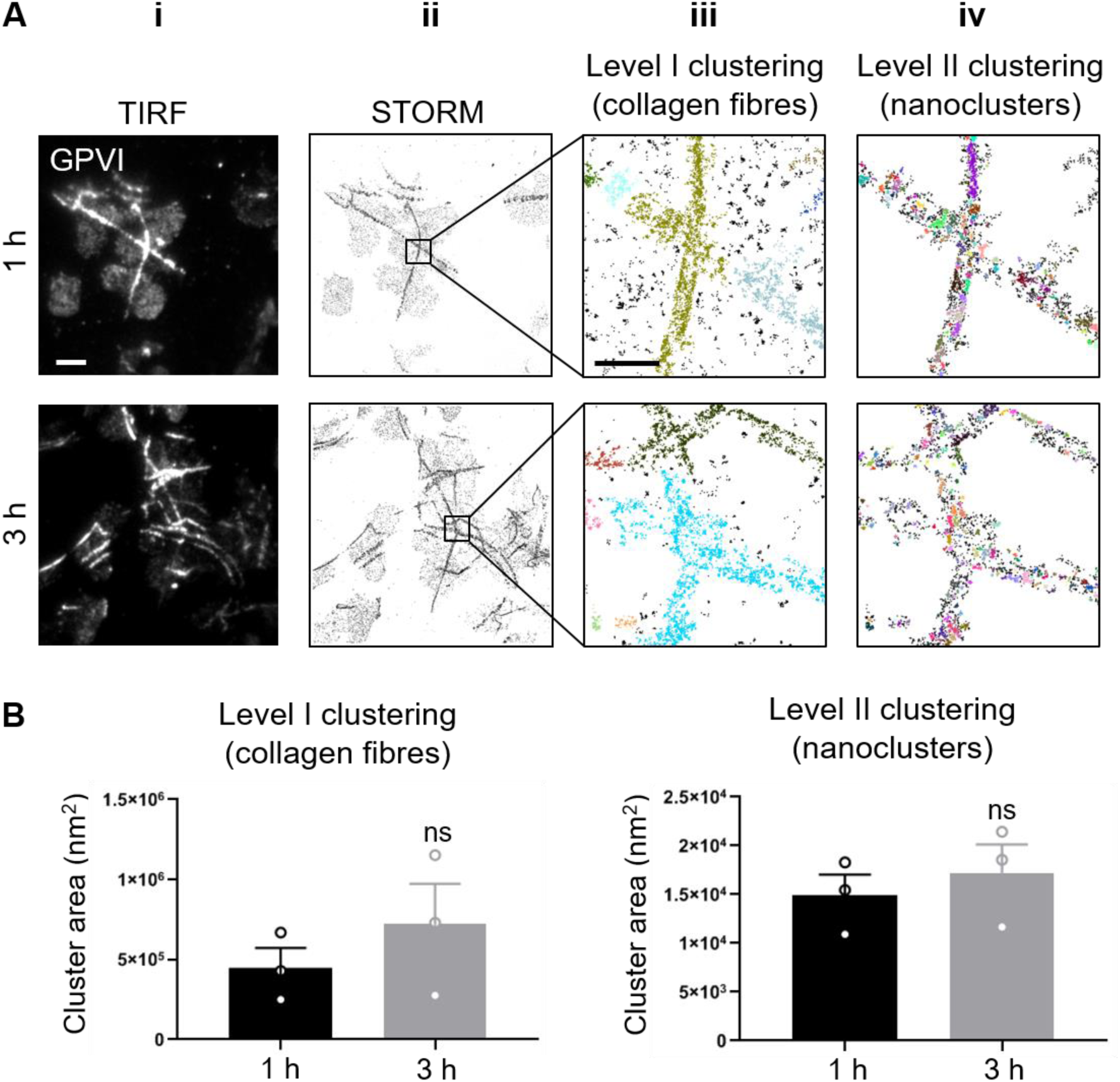
dSTORM imaging and two-level DBSCAN cluster analysis of GPVI in platelets spread on collagen for 1 h and 3 h. Washed human platelets spread on Horm collagen for 1 h and 3 h, labelled for GPVI using 1G5-Fab and imaged by TIRFM (**Ai**) and dSTORM (**Aii**). The localised data points from the dSTORM images were grouped into clusters using a two-level cluster analysis based on DBSCAN (**Aii, iii**). (**Aiii**) Level I clustering identified large clusters corresponding to collagen fibres. (**Aiv**) Level II clustering subdivided these large clusters into nanoclusters where different colours denote different clusters. (**B**) Quantitative analysis of cluster area for both Level I and Level II clustering at the different time points indicated. Bars represent mean ± SEM from three independent experiments. Significance was measured using unpaired two-tailed t-test p < 0.05: ns indicates non-significant difference. Scale bar: 5 μm (TIRF and dSTORM images) and 1 μm (cluster plots).

Cluster density and number of detections per cluster also remained constant over time (Supp. Fig. 2A, B). This suggests that GPVI-collagen interactions had stabilised within 1 h of platelet spreading and remained constant regardless of the time.

### GPVI shedding is minimal in platelets spreading on collagen

In human platelets in suspension, GPVI is cleaved upon treatment with different agonists, predominantly by ADAM10(*18*). However, whether GPVI is cleaved in platelets that have been spread on an immobilised ligand is unknown. Using an antibody raised against the cytosolic tail of GPVI, the degree of GPVI shedding in spread platelets was assessed. Intact GPVI has a molecular weight of ∼65 kDa, but when the extracellular domain is shed, a 10-kDa remnant fragment containing the cytosolic tail remains bound to the platelet. Using western blot we found that in platelets that did not contact collagen (non-adhered, NA, Fig. 4A) no GPVI remnant fragment was detected. In platelets spread on collagen, only a minimal amount of GPVI shedding was detected, with ∼15% loss of intact GPVI after 3 h (Fig. 4A, B). As a positive control, the thiol-modifying reagent *N*-ethylmalemide (NEM), which triggers activation of metalloproteinases, was added to spread platelets for 30 min before cell lysis. NEM induced ∼100% of GPVI shedding (Fig. 4A, B). We carried out two-colour immunolocalisation of the intracellular tail of GPVI (using GPVI-tail antibody) with the extracellular domain (using 1G5-Fab) to obtain spatial information on the intact receptor (Fig. 4C). Epifluorescence imaging reveals a strong overlap of two receptor domains (Fig. 4Ci, iii), which was particularly prevalent along the collagen fibres, as highlighted in the colocalisation mask (Fig. 4Civ). This indicates that this pool of GPVI has not been shed from the platelets in the 3 h timeframe, in accordance with the biochemical data.

**Fig. 4.**
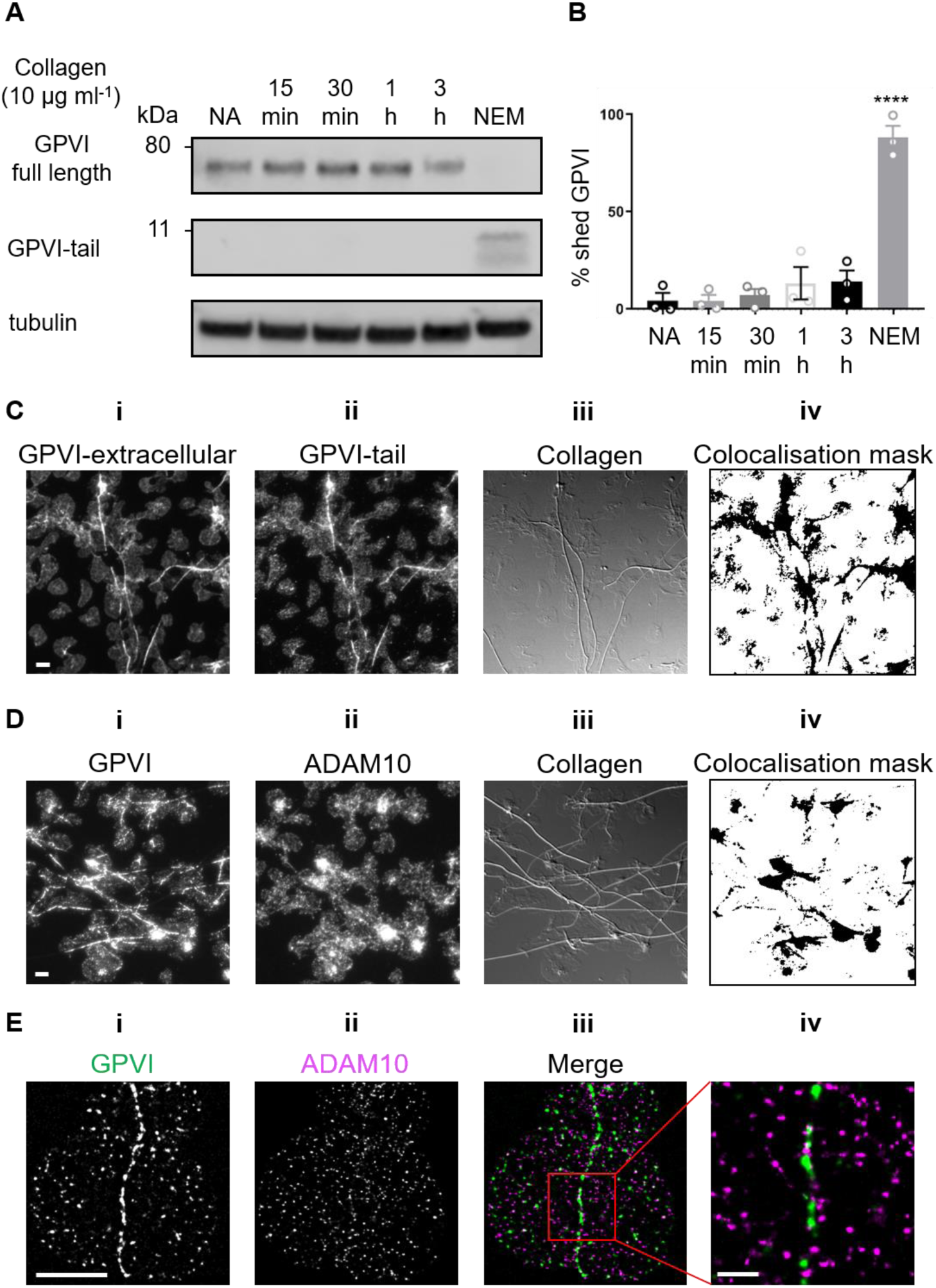
GPVI is not shed in platelets spread on fibrous collagen. (**A**) Western blot analysis of GPVI shedding using an antibody to the intracellular tail of GPVI (GPVI-tail) in washed human platelets that were either non-adhered (NA) or spread on collagen for the indicated time points. Adhered platelets treated with 2mM NEM were the positive control for shedding, tubulin was the loading control. (**B**) Quantification of the percentage of shed GPVI. Bars represent the mean ± SEM from three separate experiments. (**C**) Dual-colour epifluorescence imaging of spread platelets labelled for GPVI extracellular domain (1G5-Fab, **Ci**) and intracellular tail (GPVI-tail, **Cii**). The distribution of fibrous collagen is shown in the DIC image (**Ciii**). (**Civ**) Cololocalisation mask of the two domains of GPVI shows overlapping pixels and a high level of colocalisation along collagen fibres. (**D**) Dual-colour epifluorescence imaging of spread platelets labelled for GPVI (**Di**) and ADAM10 (**Dii**) with collagen shown in the DIC image (**Diii**). (**Div**) Colocalisation mask of GPVI and ADAM10 shows no enrichment of colocalisation at collagen fibres. (**E**) Two-colour dSTORM of GPVI (**Ei**) and ADAM10 (**Eii**). The magnified box region in the merge (**Eiii**) shows little colocalisation of GPVI and ADAM10 at a collagen fibre. Scale bar: 5 µm (whole FOVs and single platelet images) and 1 μm in (**Eiv**).

To further investigate whether GPVI shedding occurred in platelets spread on collagen we evaluated the location of the GPVI sheddase ADAM10 in relation to GPVI oligomers using two-colour epifluorescence microscopy (Fig. 4D) and dSTORM (Fig. 4E). The data demonstrate differential distribution of the two proteins, with no visible enrichment of ADAM10 at collagen fibres as was seen for GPVI (Fig. 4Di, iii, iv). Dual-colour dSTORM imaging enabled us to assess the nanoscale organization of GPVI (Fig. 4Ei) and ADAM10 (Fig. 4Eii) at the single molecule level. ADAM10 molecules rarely colocalised with the tightly packed GPVI clusters (Fig. 4Eiii, iv), suggesting that the metalloproteinase was not present in GPVI clusters, thus preventing shedding and allowing signalling to continue.

### Effect of metalloproteinases inhibition and activation on GPVI cluster formation

To investigate whether GPVI remains intact and clustered at the cell surface, dSTORM was carried out on platelets that had been spread in the presence of the broad spectrum metalloproteinase inhibitor GM6001 or the ADAM10-specific inhibitor GI254023, at concentrations previously shown to inhibit GPVI metalloproteolysis (*18, 22, 23*). If GPVI shedding did occur in spread platelets we would expect to see an increase in the measured cluster parameters when the metalloproteinase inhibitors were present. To confirm that shedding was possible, spread platelets were post-treated with NEM. Both metalloproteinase inhibitors had no effect on platelet spreading whereas NEM significantly impaired the ability of platelets to remain spread on collagen, showing a significant reduction in platelet surface area in comparison to controls (Supp. Fig. 3A, B, C). In platelets treated with either of the metalloproteinase inhibitors, discrete GPVI clusters were detected (Fig. 5A). These clusters were not significantly different from those formed in control platelets in terms of area (Fig. 5B), density (Supp. Fig. 4A) or number of detections per cluster (Supp. Fig. 4B) at both clustering levels. In contrast, very few clusters were detected in NEM-treated platelets (Fig. 5A) and there was loss of 90% of the GPVI-related single molecule detections (Fig. 5C) making the 2-level cluster analysis unfeasible. This again supports the hypothesis that GPVI shedding does not occur in platelets spread on collagen.

**Fig. 5.**
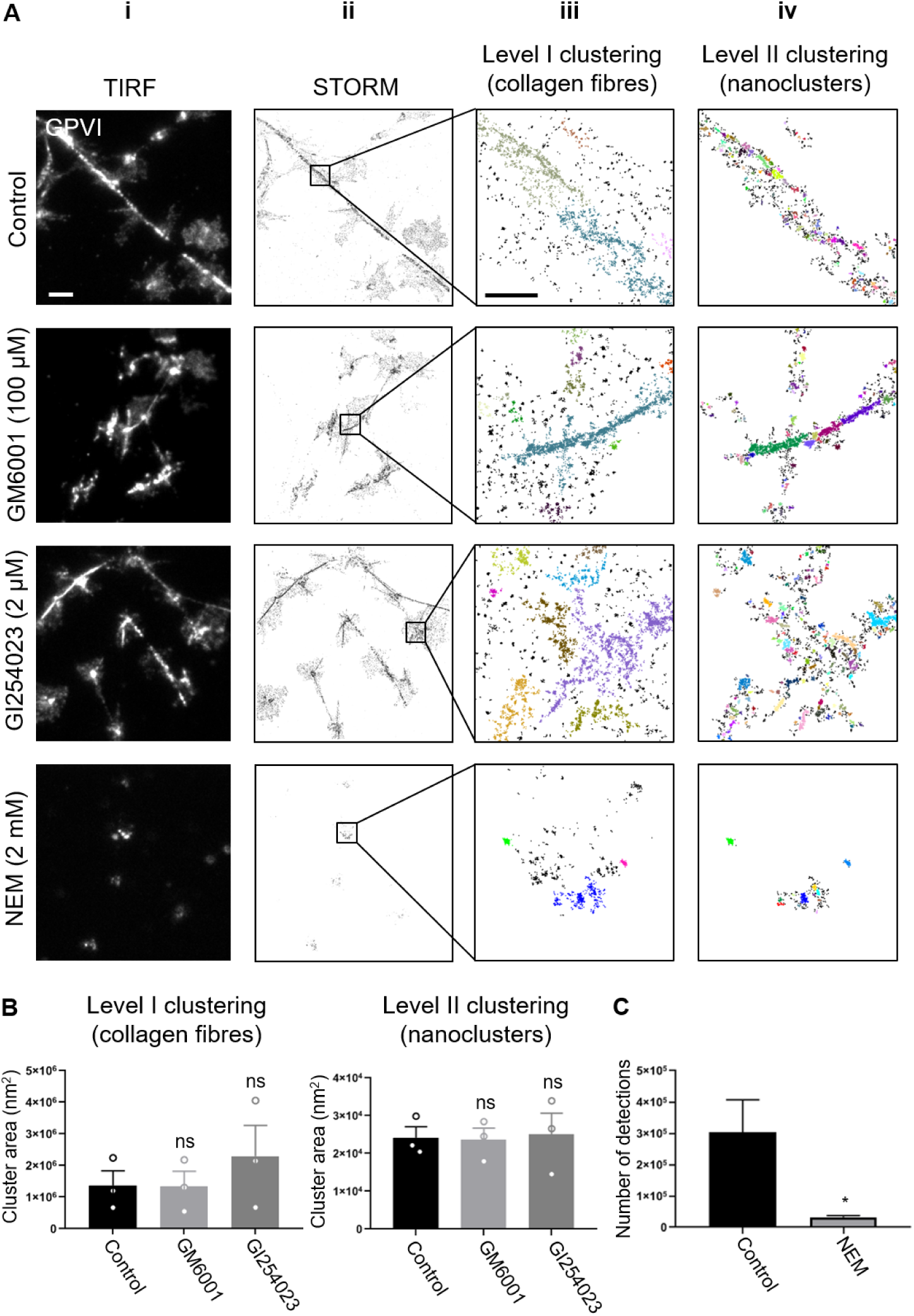
Effect of metalloproteinase inhibitors and NEM on GPVI clustering. TIRF (**Ai**) and dSTORM (**Aii**) images of GPVI in washed platelets spread on collagen for 1 h treated with DMSO (vehicle control), the metalloproteinase inhibitors GM6001 or GI254023 or NEM, which activates metalloproteinases. (**Aiii**) Level I clustering, where large clusters associated with the collagen fibres are isolated and (**Aiv**) Level II clustering where the large clusters are subdivided into nanoclusters. Different colours represent different clusters. (**B**) Quantification of the cluster size (area) at both Level I and Level II clustering. (**C**) Quantification of the number of fluorescent detections localised in dSTORM imaging of GPVI in control vs NEM-treated platelets. Bar graphs are mean ± SEM from three independent experiments. Cluster area was analysed using one-way ANOVA with Tukey’s multiple comparisons test, ns: no significant difference. Number of detections was analysed using unpaired two-tailed t-test, where * is p < 0.05. Scale bar: 5 μm (dSTORM images) and 1 μm (Level I and II cluster plots).

### Secondary mediators are not required for GPVI clustering

Release of secondary mediators, thromboxane A_2_ (TxA_2_) and adenosine diphosphate (ADP), enhance integrin activation on adherent platelets and aid thrombus development through the recruitment and activation of additional platelets(*24*). We evaluated whether secondary mediators had any effect on platelet spreading on collagen and GPVI clustering along the collagen fibres. Pre-incubation of platelets with 10 μM indomethacin and 2 U/ml apyrase, to inhibit TxA_2_ and ADP did not prevent platelet adhesion and spreading (Supp. Fig. 5A, B, C). In addition, the ability of GPVI to form clusters on collagen was not perturbed by the abolishment of secondary mediator activity (Supp. Fig. 6A, B, C, D).

### Active Src-family and Syk kinases are not required to maintain GPVI clustering on collagen

Our results show that ITAM signalling is sustained in spread platelets. In order to investigate the action of Src and Syk on the clustering of GPVI, we treated platelets with selective inhibitors of these kinases. In accordance with previous results(*17*), pre-incubation of platelets with either the Src-family kinase inhibitor PP2 or Syk kinase inhibitor PRT-060318 (PRT) significantly reduced the extent of platelet spreading on collagen (Fig. 6Aii and Supp. Fig. 7). PRT had a more marked effect with a complete abolishment of lamellipodia formation. In order to investigate whether Src-family and Syk kinases are required to maintain GPVI clustering and activation in spread platelets, we post-treated fully spread platelets for 15 min with each of the inhibitors or the vehicle control. We found that sustained spreading did not require Src-family kinase signalling as demonstrated by the lack of an effect of PP2 on platelet surface area (Fig. 6Aiii, B). However, PRT-treatment slightly reduced platelet surface area and had a visible effect on the actin cytoskeleton, with loss of stress fibres and appearance of actin nodules (Fig. 6Aiii, B). Both inhibitors impaired GPVI downstream signalling; western blot analysis showed a decrease in pPLCγ2, pSyk and pLAT levels (Fig. 6C and Supp. Fig. 8A, B, C), and Ca^2+^ imaging showed a significant reduction in the number of spiking platelets, with those spiking doing so at a much lower number, amplitude and duration than controls (Fig. 6D, Supp. movie 2). These results confirmed that Src and Syk kinases are required to maintain collagen-induced GPVI-mediated platelet activation. Finally, we evaluated the potential feedback exerted by Src and Syk kinases on GPVI receptor clustering. dSTORM imaging demonstrated that GPVI was still clustered along collagen fibres with Src-family or Syk kinases inhibition (Fig. 6E). Indeed, GPVI binding to collagen was not affected by the inhibition of either kinase activity as shown by the Level I clustering analysis (Fig. 6F and Supp. Fig. 9A). Syk inhibition perturbed the nanoscale receptor organisation within the larger collagen-associated clusters, with clusters of increased area compared to control samples (Fig. 6F). However, the density and number of detections per cluster remained unchanged (Supp. Fig. 9B).

**Fig. 6.**
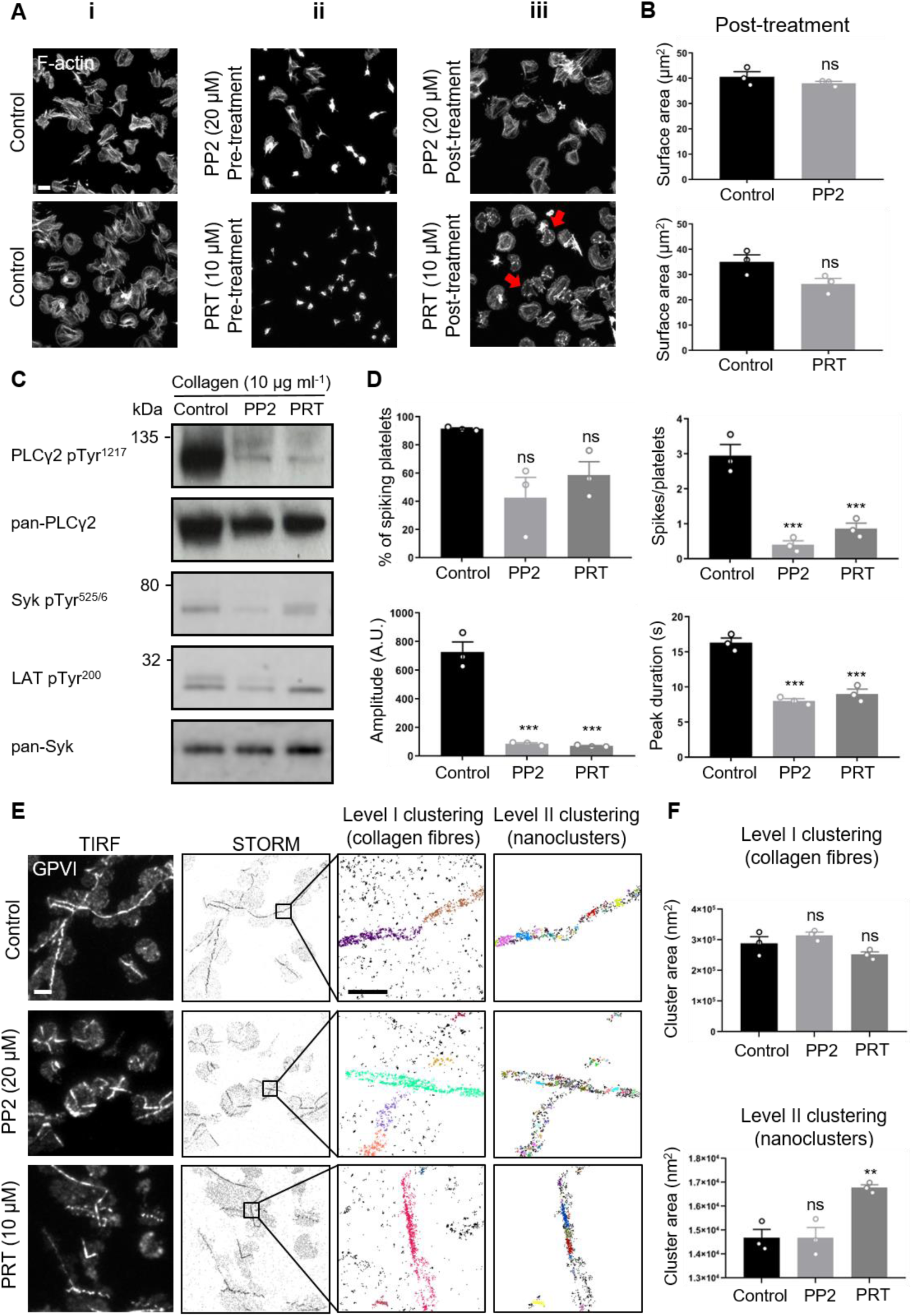
Effect of Src-family and Syk inhibitors on platelet spreading, GPVI signalling and clustering in response to fibrous collagen. (**A**) Confocal images of washed human platelets spread on collagen, labelled with phalloidin-Alexa488 to visualise F-actin. (**Ai**) Platelets pre-incubated with DMSO (vehicle control) or (**Aii**) pre-incubated with either PP2 (Src inhibitor) or PRT (Syk inhibitor) and then spread for 1h. (**Aiii**) Platelets spread for 45 min before addition of either PP2 or PRT for 15 min (post-treatment). Red arrows highlight cells where the actin cytoskeleton is disrupted. Scale bar: 5 μm. (**B**) Quantification of platelet surface area in cells treated with inhibitors post spreading. Graphs are mean ± SEM from five whole FOVs from three independent experiments. Unpaired two-tailed t-test, ns: not significantly different. (**C**) Western blot analysis of the phosphorylation state of downstream GPVI signalling proteins following post-spreading treatment with the inhibitors. (**D**) Quantification of live-cell calcium imaging of platelets labelled with Oregon green-488 BAPTA-1-AM Ca^2+^ dye, spread on collagen and then treated with vehicle control or kinase inhibitors PP2 or PRT. Bar graphs show the mean ± SEM of the percentage of spiking platelets per FOV, the number of spikes per platelets, the spike amplitude and the spike duration. In total, ∼150 platelets within a whole FOV were analysed for each individual experiment (n=3). The significance was measured using one-way ANOVA with Tukey’s multiple comparisons test (*** p < 0.001, ns: not significantly different). (**E**) TIRF and dSTORM imaging and 2-level DBSCAN cluster analysis showing the effect of PP2 and PRT post-treatment on GPVI clustering and nanoclustering. Different colours represent different clusters. (**F**) Quantification of cluster area at both Level I and Level II clustering. Graphs are mean ± SEM from n=3 experiments. Significance was measured using one-way ANOVA with Tukey’s multiple comparisons test (** p < 0.01; ns: not significantly different). Scale bar: 5 μm (TIRF and dSTORM images) and 1 μm (cluster plots).

### Inhibition of Syk but not Src kinase activity impaired clustering of integrin α2β1 along collagen fibres

To further address the effect of Src and Syk inhibition on collagen-induced platelet spreading and signalling, we investigated the distribution of the other collagen receptor integrin α2β1 using dSTORM and 2-level cluster analysis. Consistent with previous work(*17*), in spread platelets integrin α2β1 also accumulated along collagen fibres (Fig. 7A first row). Post-spreading treatment with the Src kinase inhibitor PP2 had no effect on integrin clusters (Fig. 7A second row; 7B and Supp Fig 10A, B). However, whilst post-spreading treatment with the Syk inhibitor PRT did not have any effect on cluster density or number of detections per cluster (Supp. Fig 10A, B) it resulted in a visible redistribution of α2β1 clusters (Fig 7A, third row). Large integrin α2β1 clusters were still detected (Level I analysis, Fig. 7Aiii, iv, third row) but these were no longer closely associated with the collagen, and the nanoclusters detected in the Level II analysis were significantly smaller (Fig 7B), suggesting a compromised interaction of the integrin with its ligand with Syk inhibition. These results suggest that, in contrast to GPVI, active Syk kinase is required for the maintenance of integrin α2β1 clustering along collagen fibres in spread platelets.

**Fig. 7.**
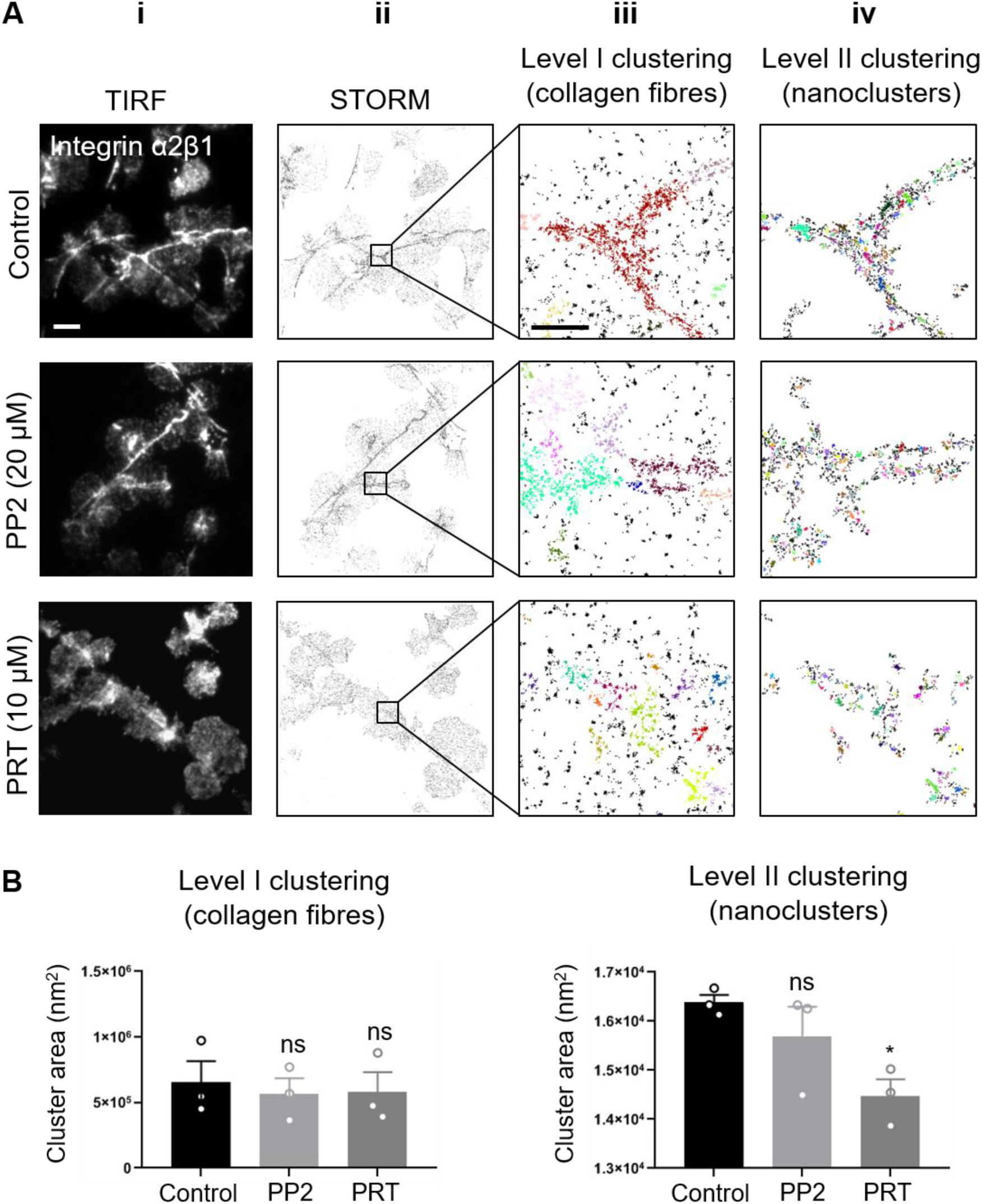
Syk but not Src-family inhibitor impairs integrin α2β1 clustering on collagen in spread platelets. (**A**) TIRF (**Ai**) and dSTORM imaging (**Aii**) and 2-level DBSCAN cluster analysis (**Aiii, iv**) of integrin α2β1 in washed human platelets spread on collagen and post-treated with DMSO (vehicle control), the Src kinase inhibitor PP2 (20 μM) or the Syk inhibitor PRT (10 μM). Different colours indicated different clusters in the cluster plots. (**B**) Quantitative analysis of cluster area. Graphs are mean ± SEM from n=3 experiments. One-way ANOVA with Tukey’s multiple comparisons test was used to evaluate the statistical significance (* p < 0.05). Scale bar: 5 μm (dSTORM images) and 1 μm (cluster plots).

## Discussion

This study has used advanced microscopy, image analysis and biochemical techniques to demonstrate that GPVI-mediated signalling is sustained in platelets adhering and spreading on an immobilised collagen surface. Our data show that GPVI clusters along the collagen fibres, leading to an enrichment of the receptor and sustained activation of downstream signalling molecules such as pSyk and pLAT and continued Ca^2+^ spiking in platelets spread for up to 3 h. Our imaging data show little overlap of GPVI with its sheddase ADAM10, which is consistent with no significant shedding of GPVI in collagen-adhered platelets. Furthermore, once bound to collagen, GPVI did not require positive feedback from Src-family and Syk kinases to maintain the clusters. This is in contrast to integrin α2β1 which requires continued Syk, but not Src, activity to remain bound to collagen fibres.

The platelet receptor GPVI has been identified as a promising antithrombotic target due to its restricted expression on platelets and megakaryocytes and the fact that loss of GPVI protects against thrombosis whilst having negligible effects on primary haemostasis (*25*). It is therefore important to understand the basic biology behind GPVI signalling and platelet activation in order to develop novel approaches to modulate GPVI receptor function. This is also important as more recent work has shown that GPVI plays a role in maintaining vascular integrity in inflammation, so this will need to be taken into account when assessing new anti-GPVI therapies(*3*).

It is known that the surface expression and distribution of GPVI is tightly regulated by a variety of stimulatory and inhibitory mechanisms to coordinate platelet responses and prevent improper activation. Indeed, GPVI spatial organisation and cluster formation at the plasma membrane play a central role in regulating signalling (*17, 26*). Platelet GPVI levels at the plasma membrane can be regulated by endocytosis(*27, 28*) or by receptor shedding, the latter being the subject of significant research(*29*). The extracellular region of GPVI can be cleaved by members of the ADAM family of sheddases(*30, 31*). ADAM10 is the major GPVI sheddase on human platelets, with ADAM17 and potentially other metalloproteinases also causing shedding of mouse GPVI(*30*). To date, GPVI shedding has been shown to be induced in platelet suspensions by a number of physiological and non-physiological stimuli, including ligands of GPVI and other platelet ITAM receptors(*32*). GPVI shedding induced by ligand engagement is thought to require receptor clustering, dissociation of calmodulin from the cytoplasmic tail and activation of the intracellular signalling cascade(*18*). Recent work using an ADAM10-specific substrate, which becomes fluorescent when cleaved, has shown that ADAM10 is constitutively active at the surface of resting platelets although GPVI is not constitutively shed(*22*). Activation of platelets with GPVI ligands such as collagen-related peptide or the purified venom protein convulxin do not alter the level of ADAM10 activity but do induce GPVI shedding, whereas exposure to pathophysiological fluid shear stress or NEM increase ADAM10 activity and cause GPVI shedding. Taken together, these observations suggest another level of control, which could include altered ADAM10 or GPVI localisation within the membrane.

This study has investigated shedding in platelets adhered to an immobilised substrate, which more closely mimics platelets adherence to collagen in vivo. We have shown that GPVI is enriched along collagen fibres and remains clustered for at least 3 h. We hypothesised that the ability of GPVI to cluster along collagen would allow signalling complexes to be formed and maintained and may safeguard GPVI from shedding by marginalisation of the sheddase ADAM10 due to steric hindrance. A similar situation is seen with the T-cell immunological synapse where the phosphatases CD148(*33*) and CD45(*34*) are marginalised from clusters of the ITAM-containing T-cell receptor, to allow signalling to be prolonged. The data support our hypothesis with minimal GPVI shedding detected and high degrees of clustered GPVI colocalising with pSyk and pLAT but very little ADAM10 and GPVI colocalisation observed. The enriched degree of colocalisation of the intra- and extracellular domains of GPVI in the dual-colour imaging combined with no evidence of remnant GPVI fragment by western blot, indicate that the receptor in contact with collagen remains intact in spread platelets. Nevertheless, GPVI shedding could be induced in spread platelets by treatment with the thiol-modifying agent NEM, commonly used to induce metalloproteinase activity. How this shedding is achieved is unclear given the distinct distributions of the two proteins.

However, NEM is not specific for ADAM10 and is membrane-permeable. NEM-treated spread platelets themselves start to detach from the collagen-coated surface. It is possible that NEM treatment could result in rearrangement of membrane proteins, enabling GPVI cleavage. Furthermore, ADAM17, as well as other platelet metalloproteinases, are known to share substrate specificity with ADAM10 and would also be activated by NEM treatment so could play a role.

Upon collagen adhesion, activated platelets release soluble agonists from the storage organelles such as ADP and TxA_2_ which act as pro-thrombotic triggers through amplification of intracellular signalling cascades, ultimately leading to enhanced integrin activation and platelet aggregation(*35*). Despite these roles in platelet responses, the discharge of secondary mediators is not needed to potentiate collagen adhesion and GPVI cluster formation, as these were not altered when platelets spread in the presence of indomethacin and apyrase. These results suggest that under these experimental conditions, secondary mediators do not function as a positive feedback loop for the initial stimulus on collagen, in contrast to their fundamental role in supporting platelet aggregation during thrombus growth. This is different to the related platelet hemITAM receptor CLEC-2, where secondary mediators play a vital role in its signalling(*36*).

Similar to GPVI, CLEC-2 also forms clusters upon interaction with its endogenous ligand podoplanin and the clustering of CLEC-2 requires active Src and Syk kinases(*37*). Since GPVI receptor cluster formation seems to maintain ITAM signalling over time, signalling may contribute to the ability of GPVI to maintain large clusters through positive feedback. Pre-incubation of platelets with the Src inhibitor PP2 or the Syk inhibitor PRT prevented complete spreading of platelets on collagen as previously shown(*17*) and therefore prevented detailed analysis of GPVI clustering. However, the effect of positive feedback of these kinases on cluster formation could be tested by addition of the inhibitor to platelets that had already undergone spreading. Treatment of spread platelets with both the inhibitors had very little effect on numbers and extent of GPVI clusters that had already formed on the collagen, with only a slight increase in the GPVI nanoclusters seen when Syk was inhibited. However, the inhibitors reduced pSyk, pLAT and pPLCγ2 and had an almost immediate effect on Ca^2+^ signalling. This demonstrates that Src-family and Syk kinases signalling are fundamental for sustaining full GPVI-mediated platelet activation. Syk, but not Src, inhibition resulted in a disruption of the actin cytoskeleton, with loss of stress fibres and a slight reduction in platelet spreading. Previous work has shown that GPVI dimer formation is dependent on the actin cytoskeleton(*17*) so the actin depolymerisation caused by Syk inhibition may disrupt GPVI dimers and could explain the appearance of slightly larger GPVI nanoclusters at the platelet surface. Syk has a well-established role in cytoskeleton organisation upon immunoreceptor stimulation in several cell types, including platelets(*38*). Work in B-cells has also shown a differential dependence upon Src and Syk family kinases in B-cell receptor (BCR) signalling. Under conditions of BCR clustering by multivalent ligands, Syk is sufficient to induce signalling. However, with monomeric ligands where no clustering is initiated, both Syk and Src family kinases are required for BCR activation(*39*). In spread platelets, actin disruption caused by inhibiting Syk may be mediated through the action of Syk on the Vav-family of Rho/Rac GTPase guanine nucleotide exchange factors (GEFs), which have been shown to require Syk-mediated tyrosine phosphorylation to become activated(*40*). Vav1 and Vav3 play redundant roles in platelet activation by collagen and loss of these proteins prevents PLCγ2 phosphorylation and full platelet spreading(*41*). Together, these results suggest that, whereas sustained collagen-induced platelet activation requires intact Src and Syk activity, prolonged GPVI cluster formation does not.

In platelets there is a second receptor for collagen, integrin α2β1, which has previously been shown to also cluster along collagen fibres(*17*). In contrast to GPVI, our current and previous(*42*) results show that Syk, but not Src, kinase activity is required for maintaining integrin α2β1 clustering along collagen. Unlike GPVI, integrins fluctuate between an ‘on’ and ‘off’ state and only the conversion into an active conformation allows high-affinity ligand binding(*43, 44*). The F-actin cytoskeleton plays an important role in the up-regulation of integrin activity(*45*) through the engagement of different adaptor proteins such as kindlin, vinculin and talin which couple integrins to the F-actin(*46*). Indeed, the importance of this interaction has been demonstrated by platelet-specific talin1 knockout mice, where loss of talin from platelets caused a massive reduction in β1 integrin-mediated platelet adhesion under flow conditions, with no stable adhesion or microthrombi formed(*47*). Our results suggest that the actin disruption caused by Syk-inhibition causes inactivation of the integrin and its subsequent dissociation from collagen. In contrast, as GPVI clusters are largely unaffected by PRT treatment it implies that, once bound to collagen, GPVI remains bound and does not have such a dependence on the actin cytoskeleton. As integrin α2β1 is the main adhesive receptor, dissociation from collagen would have a negative impact on platelet adhesion, especially under flow conditions where GPVI has been shown to be the main signalling receptor and integrin α2β1 plays a role in stable adhesion(*48*).

Taken together our results support a model in which GPVI clustering induced by immobilised collagen triggers the assembly of signalling complexes at the platelet surface via the engagement of downstream signalling proteins, increasing the avidity for ligand-binding and strengthening the signal. We show that GPVI clustering is maintained over time and helps to prevent negative regulation mediated by the sheddase ADAM10. The sustained, primarily Syk-mediated, signalling is required to prolong calcium signals, maintain actin cytoskeletal arrangements and integrin α2β1 engagement with collagen to ensure that platelets remain properly spread and adhered to the substrate. Thus, GPVI clustering at collagen is important for initiation of a stable thrombus and the prevention of bleeding associated with disrupted vascular integrity driven by inflammation, where GPVI plays a critical role(*3*). Due to the relatively minor role of GPVI in haemostasis, this receptor has been identified as a promising anti-thrombotic target. The selective inhibition of receptor clustering induced by collagen provides a new avenue for the development of novel therapeutic approaches to improve the functional outcomes of a variety of thrombotic and inflammatory conditions.

## Materials and methods

### Reagents and antibodies

GPVI and the GPVI-cytoplasmic tail were detected using 1G5 Fab and affinity-purified anti-GPVI cytoplasmic tail IgG(*49*), respectively. The other antibodies used in the study were purchased from commercial suppliers. Monoclonal antibodies: anti-CD49b (integrin α2β1, abD Serotec, Oxford, UK); anti-ADAM10 [clone 11G2] (Abcam, Cambridge UK); anti-phosphotyrosine (clone 4G10). Polyclonal antibodies: anti-LAT IgG (Millipore Merck, Abingdon, UK); anti-Syk IgG (#sc1077) and anti-PLCγ2 IgG (Q-20):sc-407 (Santa Cruz Biotechnology, Dallas, USA); anti-phospho LAT Tyr^200^ (Abcam, Cambridge UK); anti-phospho Syk Tyr^525/526^, anti-phospho PLCγ2 Tyr^1217^ and AlexaFluor 647-conjugated anti-phospho-Syk Tyr^525/526^ (Cell Signaling Technology, Hitchin, UK). All the fluorescent secondary antibodies, phalloidin-Alexa 488 and Oregon green 488 BAPTA-1-AM Ca^2+^ dye used for imaging, were purchased from Thermo Fisher Scientific (Waltham, MA, USA). The HRP- and fluorescence-conjugated secondary antibodies, used for immunoblotting, were obtained from Amersham Biosciences (GE Healthcare, Bucks, UK) and antibodies-online.com GmbH (Aachen, Germany), respectively. Horm collagen was obtained from Nycomed Pharma GmbH (Munich, Germany). 10% neutral buffered Formalin solution and Hydromount solution were purchased from Sigma (Poole, UK) and National Diagnostics (Atlanta, USA), respectively. The broad spectrum metalloproteinases inhibitor GM6001 and N-ethylmaleimide (NEM) were obtained from Millipore Merck (Abingdon, UK). The ADAM10 inhibitor GI254023 was purchased from Scientific Laboratory Supplies (Nottingham, UK). The Src-family kinases inhibitor PP2 and Syk inhibitor PRT-060318 were purchased from TOCRIS (Abingdon, UK) and Caltag Medsystems (Buckingham, UK), respectively. The inhibitors indomethacin and apyrase were obtained from Sigma (Poole, UK).

### Human platelet preparation

Human washed platelets were prepared from blood samples donated by healthy, consenting volunteers under the following licence: ERN_11-0175 ‘The regulation of activation of platelets’ (University of Birmingham). Blood was drawn via venipuncture into the anticoagulant sodium citrate and then acid/citrate/dextrose (ACD) added to 10% (v:v). Blood was centrifuged at 200 *× g* for 20 min. Platelet-rich plasma (PRP) was collected and centrifuged at 1,000 *× g* for 10 min in the presence of 0.1 μg ml^-1^ prostacyclin. Plasma was removed and the platelet pellet was resuspended in modified Tyrode’s buffer (129 mM NaCl, 0.34 mM Na_2_HPO_4_, 2.9 mM KCl, 12 mM NaHCO_3_, 20 mM HEPES, 5 mM glucose, 1 mM MgCl_2_; pH 7.3) containing ACD and 0.1 μg ml-1 prostacyclin before centrifugation at 1,000 *× g* for 10 min. The washed platelet pellet was resuspended in modified Tyrode’s buffer, left to rest for 30 min at room temperature (RT) and the platelet count adjusted to the desired concentration.

### Platelet spreading and staining

For confocal and dSTORM imaging, 13 mm #1.5 glass coverslips (VWK, UK) and 35 mm #1.5 (0.17 mm) uncoated glass-bottomed dishes (MatTek Corporation, Ashland, MA, USA) were coated with 10 μg ml^-1^ Horm collagen diluted in manufacturer-supplied diluent, and left overnight at 4°C, before being blocked in 5 mg ml^-1^ BSA for 1 hour at RT. Washed and rested platelets were diluted to 2×10^7^ cells ml^-1^ in modified Tyrode’s buffer and allowed to spread on the collagen-coated surface for the required times at 37 °C. When required, washed platelets were labelled with 2 μg ml^-1^ 1G5-Fab against intact GPVI for 10 min at 37 °C prior to spreading. Where stated, platelets were also incubated with specific inhibitors before and/or after spreading on collagen. Adhered cells were PBS-washed, fixed for 10 min with 10% neutral buffered Formalin solution, permeabilized for 5 min with 0.1% (v:v) Triton X-100 in PBS, blocked with 1% (w/v) BSA + 2% (v:v) goat serum in PBS and then labelled with antibodies or phalloidin for 1 h at RT as required. For confocal and epifluorescence imaging, samples were mounted on glass slides using Hydromount solution and stored at RT. Samples for direct stochastic optical reconstruction microscopy (dSTORM) were stored in PBS at RT until imaged.

### Confocal imaging

Spread platelets mounted on glass slides were imaged using a Leica TCS SP2 confocal microscope and the 63x 1.4NA objective. For each condition, 5 z-stacks (step size 0.25 µm) were imaged at random positions on the coverslip at a resolution of 1024 ⨯ 1024 pixels and a zoom of 4. Phalloidin-Alexa 488-labelled platelets were imaged with a 488 argon laser and Alexa-647 labelled platelets with a 647 HeNe laser. Dual-colour labelled platelets were imaged simultaneously. Single-plane reflection images were also taken to monitor the distribution of the collagen fibrils. The images were then processed using ImageJ v1.48 (NIH, Bethesda, USA).

### Colocalisation analyses

Qualitative and quantitative colocalisation analyses of GPVI and phospho-proteins were performed using Fiji v1.52(*50*). For the qualitative colocalisation analysis, the maximum intensity projections for each channel were thresholded using the ImageJ Default method (a variation of the IsoData algorithm) and multiplied together to obtain colocalisation masks showing only the colocalising pixels. For the quantitative colocalisation analysis, regions of interest were drawn around single platelets or groups of platelets randomly chosen within the field of view (FOV) and the Plugin Coloc2 was used to measure the degree of colocalisation through the Pearson’s correlation coefficient.

### Epifluorescence imaging

Dual-colour imaging of Alexa 488 and 647-labelled platelets spread on collagen was performed using a Zeiss Axio Observer 7 Epifluorescent microscope equipped with a 63x 1.4NA oil immersion lens, Colibri 7 LED light source, Zeiss Filter sets 38 and 50 for GFP/FITC and Cy5/647, respectively and Hammamatsu ORCA Flash 4 LT sCMOS camera for image acquisition. DIC images were also taken (intensity= 7.0V and exposure time= 100 ms) to visualise the collagen distribution. For each condition, 5 random FOVs were imaged (intensity= 20% and 25% and exposure time: 200 ms and 100 ms for 488 and 647, respectively) and then analysed in Fiji v1.52(*50*).

### Platelet spreading analysis

The confocal and epifluorescence images of platelets spread on collagen were assessed for platelet count and surface area using specalized semi-automated workflow(*51*) built on the open-source KNIME software(*52*) and Ilastik(*53*). A pixel classifier was used to construct a binary cell segmentation and the centre of each single platelet was then set manually to facilitate the separation of touching cells. The coordinates of the cell centre positions were then used to generate the final segmentation by employing a watershed-based transformation algorithm. Only objects with a size > 1 μm^2^ were included in the final per cell measurements including area and circularity.

### STORM imaging

Super-resolution imaging of GPVI receptors was performed using a Nikon N-STORM system in TIRF and dSTORM mode with a 100x 1.49NA TIRF objective lens. The microscope system includes a Ti-E stand with Perfect Focus, an Agilent Ultra High Power Dual Output Laser bed with 170-mW 647-nm and 20-mW 405-nm lasers for the fluorophore excitation and an Andor IXON Ultra 897 EMCCD camera for the image capture. DIC and TIRF images were acquired on random FOVs containing both platelets and collagen fibres. An oxidizing and reducing buffer (100 mM MEA, 50 μg ml^-1^ glucose oxidase and 1 μg ml^-1^ catalase diluted in PBS, pH 7.5(*54*)) was used to allow the photoswitching performance. For single colour (Alexa647) imaging of labelled GPVI, the N-STORM emission cube was used, combined with the gradual increase of the 405 laser power (5% every 30 sec) to reactivate the fluorophore switching. Simultaneous dual-colour (Alexa647 and Alexa488) imaging was carried out using the Nikon Quad Cube. For each FOV, 20,000 frames were acquired in Nikon NIS Elements v4.5 software using an exposure time of 20 ms, gain 300 and conversion gain 3. To generate the final super-resolved images, the ThunderSTORM plugin for Fiji was implemented with the application of the Gaussian PSF model and maximum likelihood fitting(*55*). Drift correction and photon intensity filtering (>1000 photons) were applied within ThunderSTORM to post-process the reconstructed images. To minimise the ‘multiblinking’ artifacts, detections within 75 nm of another detection, either in the same or subsequent frames, were merged. The normalised Gaussian method was then used to allow visualisation of the reconstructed STORM images which contain the spatial coordinates for each detected fluorescent blink. The image datasets were then exported as text files and analysed for clustering in KNIME software(*52*).

### Two-level cluster analysis of dSTORM data

Two-level cluster analysis was performed by first segmenting large clusters, corresponding to collagen fibres (Level I), and then segmenting denser clusters within the large clusters using different algorithm parameters (Level II). This analysis was implemented using the R library RSMLM(*42*) and KNIME(*52*) (workflow available upon request). Both segmentation levels were calculated using density-based spatial clustering of applications with noise (DBSCAN)(*21*). The radii of the local neighbourhood were set to 75nm and 30nm for Level I and II respectively, and the minimum number of reachable detections was set to either 3 or 10 for the different levels. Clusters with less than either 100 (Level I) or 10 (Level II) detections were removed. The union of circular regions (radius = 30 nm) positioned around each point in a cluster was used to measure the cluster area applying a grid of 5 nm pixels and image-based dilation. The analysis was performed on images using the whole FOV and the quantitative data relative to all clusters were outputted to an Excel spreadsheet.

### Measurements of Ca^2+^ mobilisation

For Ca^2+^ mobilisation analysis, washed platelets, diluted to 2 × 10^8^ cells ml^-1^ in modified Tyrode’s buffer, were incubated for 45 min at 37°C with 1 μM Oregon green-488 BAPTA-1-AM and centrifuged at 1000 × *g* for 10 min in the presence of 2.8 μM PGI_2_ and ACD. The platelet pellet was resuspended in the same volume of modified Tyrode’s buffer and rested for at least 30 min before diluting to 2 × 10^7^ platelets ml^-1^ for spreading. Rested platelets were then plated on collagen-coated MatTek dishes, prepared as described above, to adhere and spread. Live-cell imaging of Ca^2+^ mobilisation was tracked using the Zeiss Axio Observer 7 Epifluorescent microscope as detailed above. Where indicated, the Src inhibitor PP2 (20 µM) and the Syk inhibitor PRT-060318 (10 µM) or the vehicle (DMSO) were added to the platelets after 45 min of spreading. Images were then acquired every 1 sec for 2 min using Zen Pro v2.3 software. Once processed in Fiji v1.52(*50*), the percentage of spiking platelets, number of spikes per platelet, amplitude and peak duration were measured using MATLAB (Mathworks, Inc., Natick, MA) within one representative FOV (∼150 cells) for each condition. Signals were baseline-corrected by application of a linear top-hat filter. A threshold of 20 fluorescence units was set for identification of spikes and the duration of a spike was defined as the time for fluorescence to reduce to below this threshold.

### Protein phosphorylation and receptor shedding assay

Washed platelets were diluted to 5 × 10^8^ cells ml^-1^ in modified Tyrode’s buffer supplemented with 2 mM CaCl_2_ and allowed to spread on a collagen-coated surface (10 μg ml^-1^) for specific time periods at 37 °C. Non-adherent platelets were removed and lysed in 2x lysis buffer (300 mM NaCl, 20 mM Tris, 2 mM EGTA, 2 mM EDTA, and 2% NP-40 detergent, pH 7.4 supplemented with protease inhibitors: 2 mM Na3VO4, 2 mM 4-(2-Aminoethyl) benzenesulfonyl fluoride hydrochloride (AEBSF), 10 μg ml^-1^ leupeptin, 10 μg ml^-1^ aprotinin and 1 μg ml^-1^ pepstatin). Adherent and spread platelets were PBS-washed and then lysed in 1x lysis buffer. The protein concentration of both lysates was measured using a Bio-Rad Protein Assay, diluted to the same concentration in lysis buffer and resuspended in appropriate volumes of 5x reducing sodium dodecyl sulfate (SDS) sample buffer (10 mg ml^-1^ SDS, 25% (v:v) 2-mercaptoethanol, 50% (v:v) glycerol, 25% (v:v) Stacking buffer (0.5 M Tris HCl, pH 6.8), containing a trace amount of Brilliant Blue) and stored at −20°C.

### Western blot

Lysate samples were denatured at 95°C for 10 min and centrifuged at 18000 *× g* for 5 min at 4°C to pellet insoluble material. Pre-cast polyacrylamide gels, Bolt 4-12% Bis-Tris Plus (Invitrogen) were used to run samples. Resolved proteins were transferred to PVDF membrane (Trans-Blot Turbo RTA transfer kit, LF PVDF from Bio-Rad). After transfer, the membrane was blocked for 1 h in 5% (w:v) BSA buffer and then incubated with specific primary antibodies overnight at 4°C. Membrane washing (3 × 10 min) to remove unbound primary antibodies was executed in TBST (50 mM Tris-Cl pH 7.5, 150 mM NaCl, 0.05% (v:v) Tween 20) before incubating with fluorophore- or HRP-conjugated secondary antibodies (1:10,000) for 1 h at RT. To remove unbound secondary antibodies, washing (3 × 10 min) was repeated. For chemiluminescence-based detection of the proteins, the membrane was then incubated with the Pierce ECL Western Blotting Substrate (Thermo Fisher Scientific). Both chemiluminescent and fluorescent antibody binding was visualised using the Odyssey Fc System (LI-COR). Quantification of the intensity of the bands was performed using Image Studio v5.2. The percentage of cleaved GPVI was calculated using the band intensities of full-length GPVI (∼62kDa) and GPVI tail remnant (∼10kDa) as follows: % shed GPVI = (remnant GPVI / (full-length GPVI + remnant GPVI)) * 100.

### Statistical Analysis

Results are expressed as means ± standard error of the mean (SEM). Data were analysed using GraphPad Prism 8 software (GraphPad Software, Inc., La Jolla, CA, USA). Significant differences were determined using unpaired two-tailed Student’s *t*-test or one-way Analysis of variance (ANOVA) with Tukey’s post-hoc test for multiple comparisons. The significance was set at *p* ≤ 0.05.

## Supplementary Materials

**Supp. Fig. 1**. Western blot quantitative analysis of phosphorylated Syk and LAT from lysates of platelets spread on collagen for different time points.

**Supp. Fig. 2**. Two-level DBSCAN cluster analysis of GPVI in platelets spread on collagen for 1 h and 3 h.

**Supp. Fig. 3**. Effect of metalloproteinase inhibitors and NEM on platelet spreading.

**Supp. Fig. 4**. Two-level DBSCAN cluster analysis of GPVI in platelets spread on collagen in the presence of metalloproteinase inhibitors.

**Supp. Fig. 5**. Platelet spreading on collagen occurs independently of secondary mediators signalling.

**Supp. Fig. 6**. Inhibition of secondary mediators does not perturb GPVI cluster formation in platelets spread on collagen.

**Supp. Fig. 7**. Effect of Src-family and Syk kinases of platelet spreading and GPVI clustering on collagen.

**Supp. Fig. 8**. Western blot quantitative analysis of phosphorylated PLCγ2, Syk and LAT from lysates of platelets spread on collagen and and post-treated with Src-family and Syk inhibitors.

**Supp. Fig. 9**. Two-level DBSCAN cluster analysis of GPVI in platelets spread on collagen and post-treated with Src-family and Syk inhibitors.

**Supp. Fig. 10**. Two-level DBSCAN cluster analysis of integrin α2β1 in platelets spread on collagen and post-treated with Src-family and Syk inhibitors.

**Supplementary Movie 1**. Calcium mobilisation in platelets spread on collagen for different time periods.

**Supplementary Movie 2**. Calcium mobilisation in spread platelets after Src or Syk kinase inhibition.

## Acknowledgements

We thank Deirdre Kavanagh, the COMPARE microscope officer, for support with imaging and Mark Probert for preliminary imaging data. We also thank Steve Thomas, Abdullah Khan, Julie Rayes and Christopher Smith for helpful discussions.

## Author contributions

C.P. did experimental work, analysed the data, prepared figures and wrote the manuscript. J.A.P. implemented the cluster analysis and platelet spreading workflows. C.O’S. created the calcium analysis code and analysed the data. R.K.A. and E.E.G. provided key reagents and expertise. S.P.W. provided supervision, devised experiments and interpreted the data. N.S.P. conceived the project, devised experiments, interpreted the data, prepared figures and wrote the manuscript. All authors reviewed the manuscript.

