## Supplemental Figures for "Immobilised collagen prevents shedding and induces sustained GPVI clustering and signalling in platelets"

#### Supplementary Fig. 1

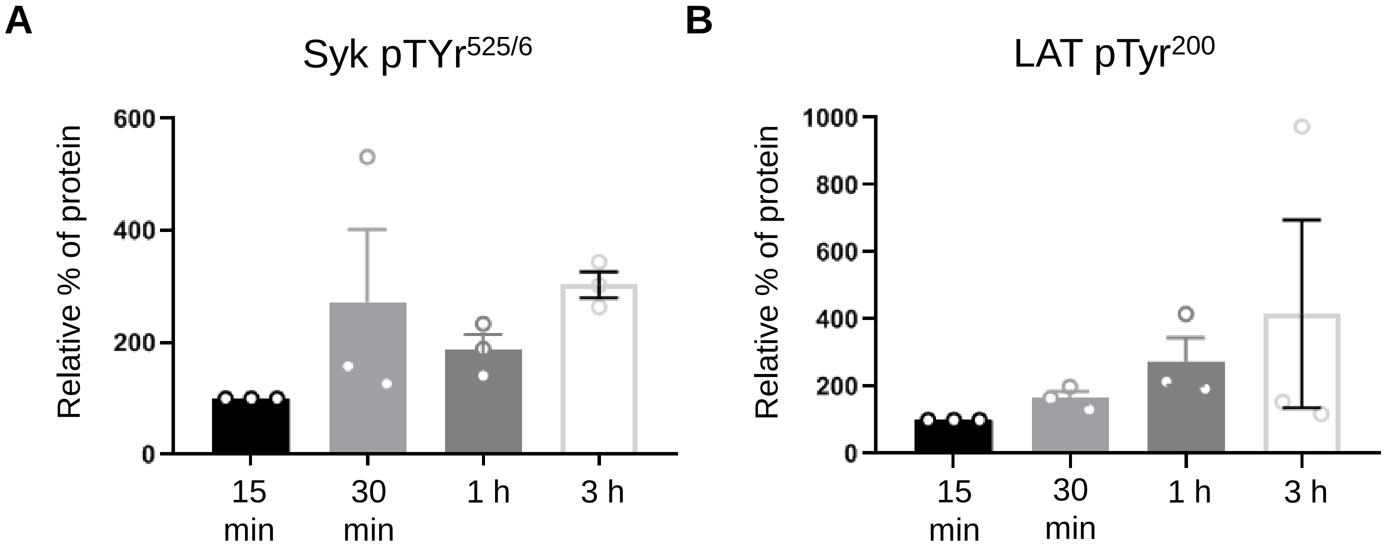


**Supp. Fig. 1 Western blot quantitative analysis of phosphorylated Syk and LAT from lysates of platelets spread on collagen for different time points.**
Quantification of phospho-Syk (**A**) and phospho-LAT (**B**) levels, normalised against the pan signals from lysates of platelets spread on collagen for the indicated time points. Data are reported as mean ± SEM relative % of protein (15 min set to 100%) from three independent experiments. No significant difference between time points was detected using one-way ANOVA with Tukey's multiple comparisons test.

**Supplementary Fig. 2**

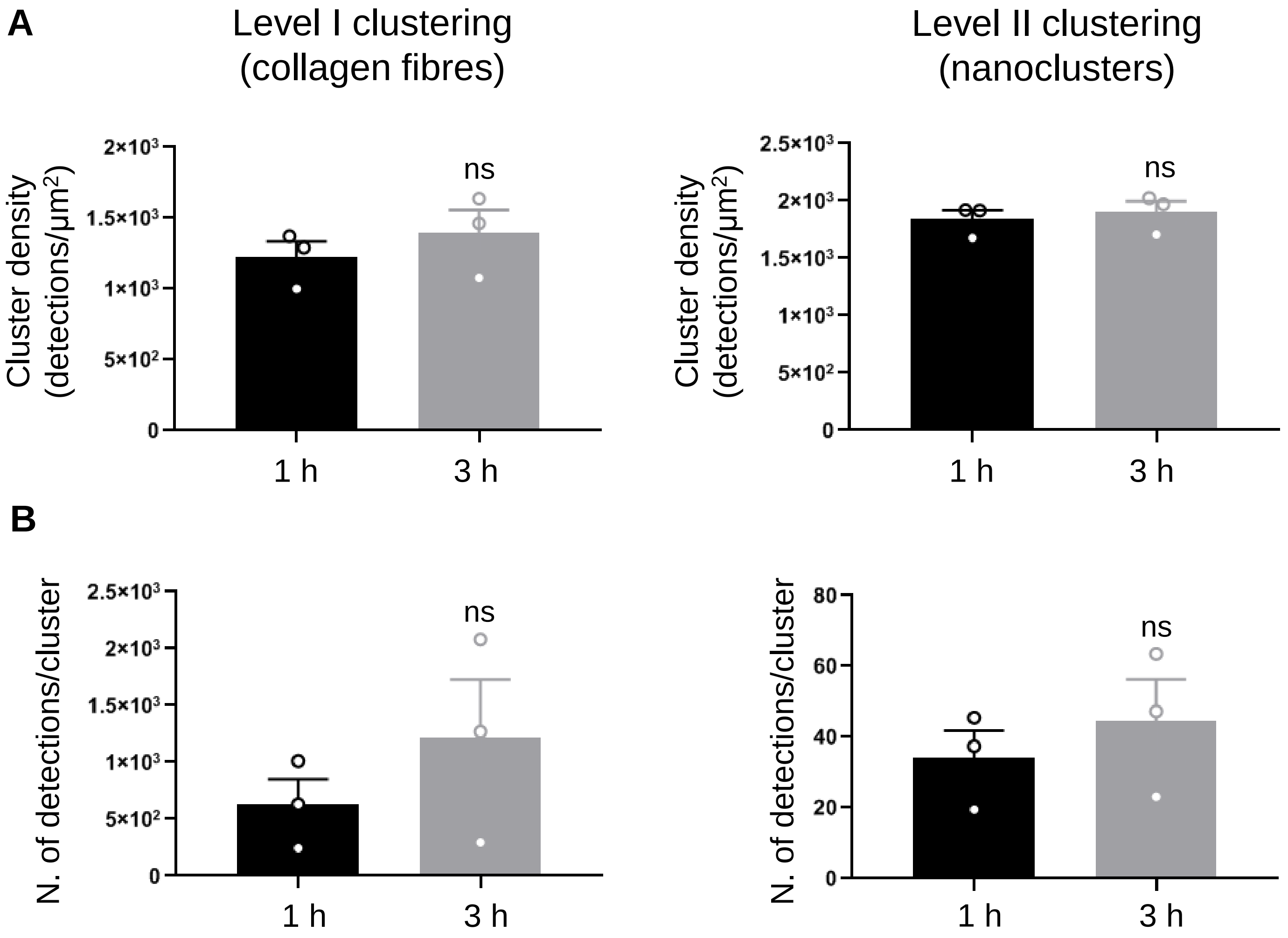


**Supp. Fig. 2 Two-level DBSCAN cluster analysis of GPVI in platelets spread on collagen for 1 h and 3 h.** Two-level DBSCAN cluster analysis of dSTORM images labelled for GPVI in platelets spread on collagen for 1 h and 3 h. Quantitative analysis of GPVI cluster density (**A**) and number of detections per cluster (**B**) for both Level I and Level II clustering for the indicated time points. Bars represent mean ± SEM from three independent experiments. Significance was measured using unpaired two-tailed t-test, $p < 0.05$: ns indicates no significant difference.

#### Supplementary Fig. 3

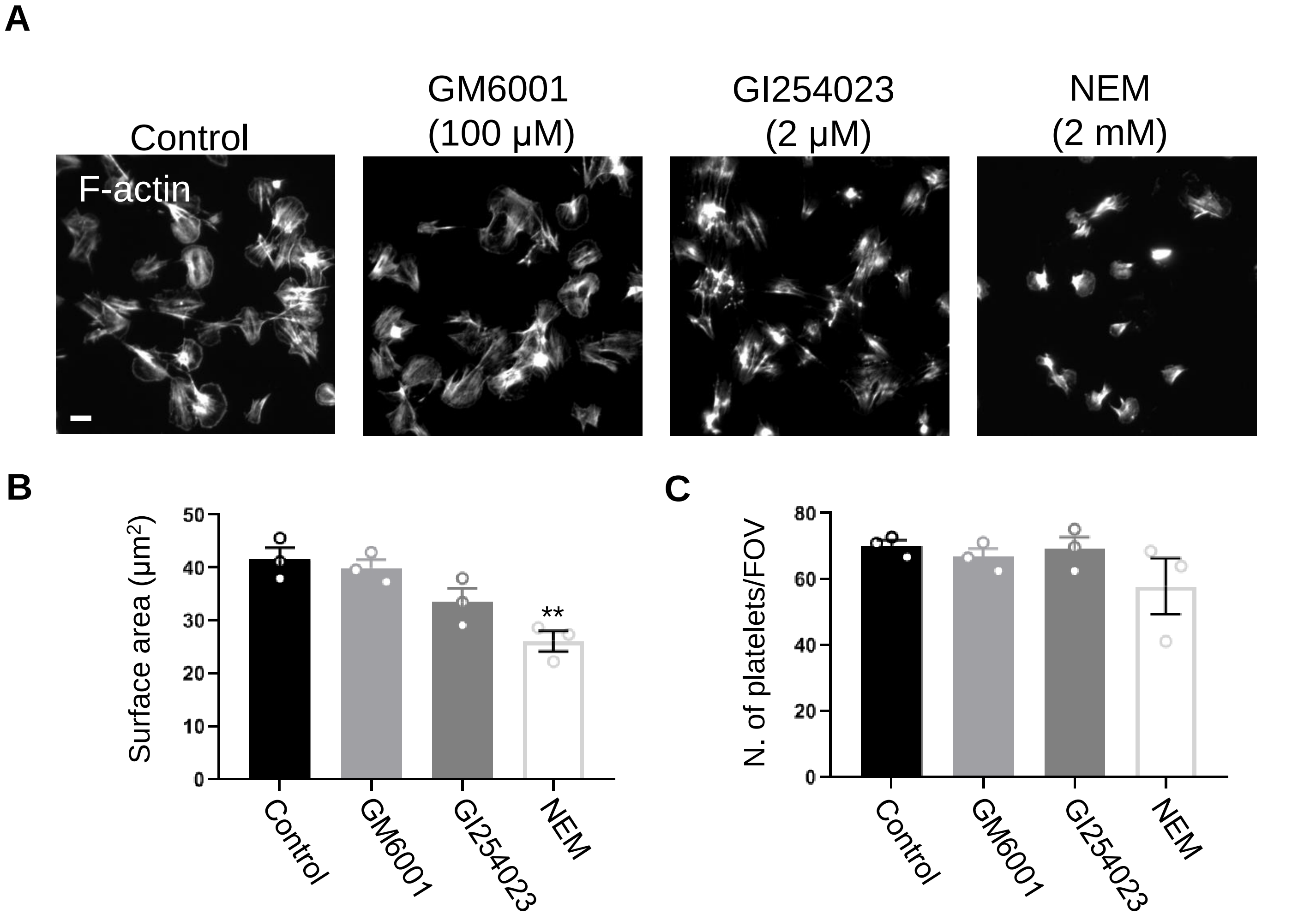


**Supp. Fig. 3 Effect of metalloproteinase inhibitors and NEM on platelet spreading.** Epifluorescence microscopy imaging of washed human platelets spread on collagen for 1h treated with DMSO (vehicle control), the metalloproteinase inhibitors GM6001 or GI254023 or the activator of metalloproteinases NEM and labelled for F-actin (**A**). Quantification of platelet surface area (**B**) and platelet number per field of view (**C**) in control and treated platelets from five FOVs from three independent experiments. Data are expressed as mean ± SEM. Significance was calculated using one-way ANOVA with Tukey's multiple comparisons test (** $p < 0.01$). Scale bar: 5 µm.

**Supplementary Fig. 4**

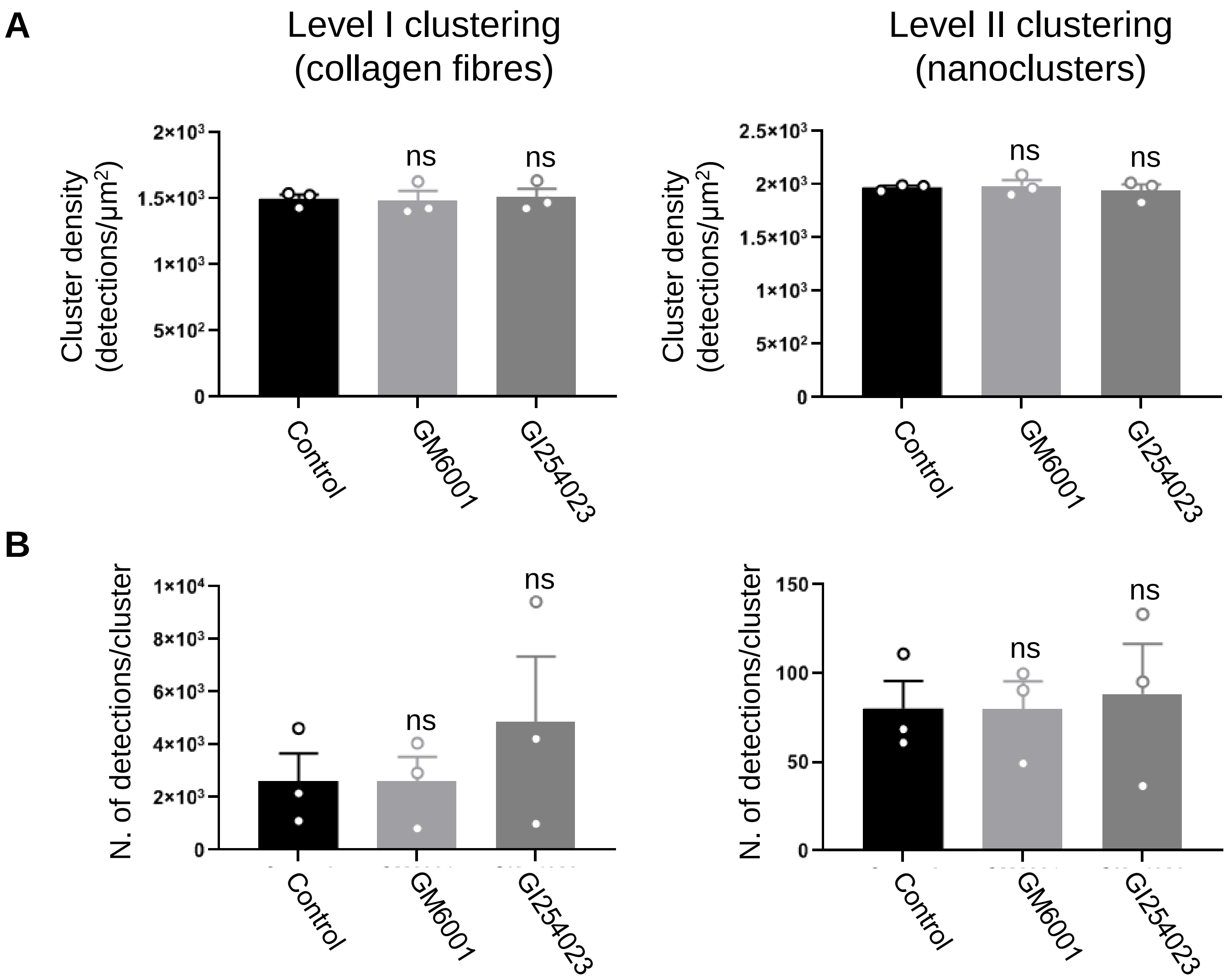


**Supp. Fig. 4 Two-level DBSCAN cluster analysis of GPVI in platelets spread on collagen in the presence of metalloproteinase inhibitors.**
Two-level DBSCAN cluster analysis of dSTORM images labelled for GPVI in platelets spread on collagen for 1 h treated with DMSO (vehicle control) or the metalloproteinase inhibitors GM6001 (100 µM) or GI254023 (2 µM). Quantitative analysis of GPVI cluster density (**A**) and number of detections per cluster (**B**) for both clustering levels in control and treated platelets. Data are reported as mean ± SEM from three independent experiments. One-way ANOVA with Tukey's multiple comparisons test ($p < 0.05$) shows non-significant (ns) difference in the clustering parameters indicated between the control and platelets treated with metalloproteinase inhibitors.

#### Supplementary Fig. 5

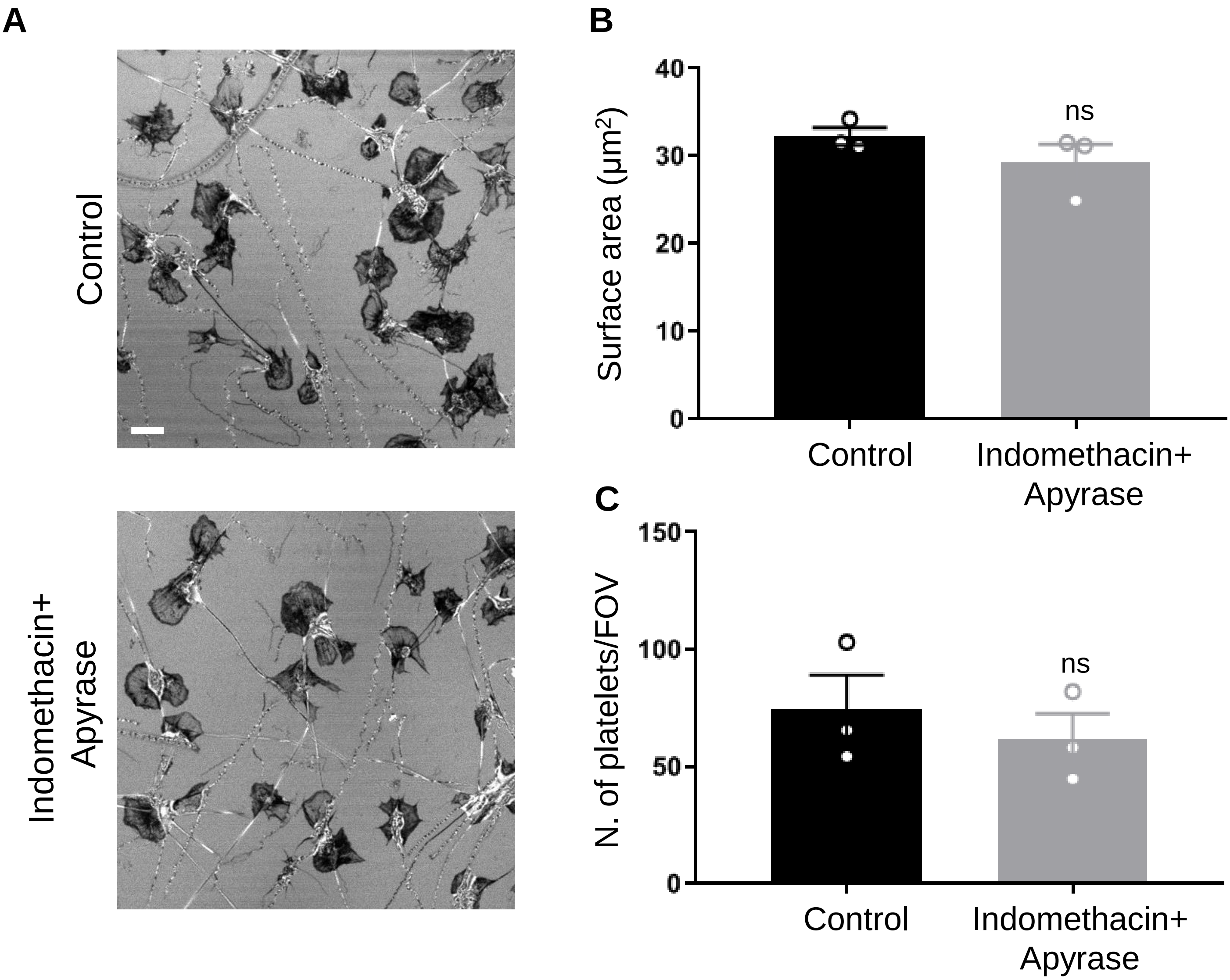


**Supp. Fig. 5 Platelet spreading on collagen occurs independently of secondary mediators signalling.** Confocal reflection images of washed human platelets spread on collagen for 1 h in the presence of DMSO (vehicle control) or 10 μM indomethacin and 2 U/ml apyrase (**A**). Measurement of platelet surface area (**B**) and platelet number per field of view (**C**) in control and platelets treated with secondary mediator inhibitors from five FOVs. Graphs are mean ± SEM from three independent experiments. Significance was measured using unpaired two-tailed t-test with $p < 0.05$: ns indicates no significant difference. Scale bar: 5 μm.

**Supplementary Fig. 6**

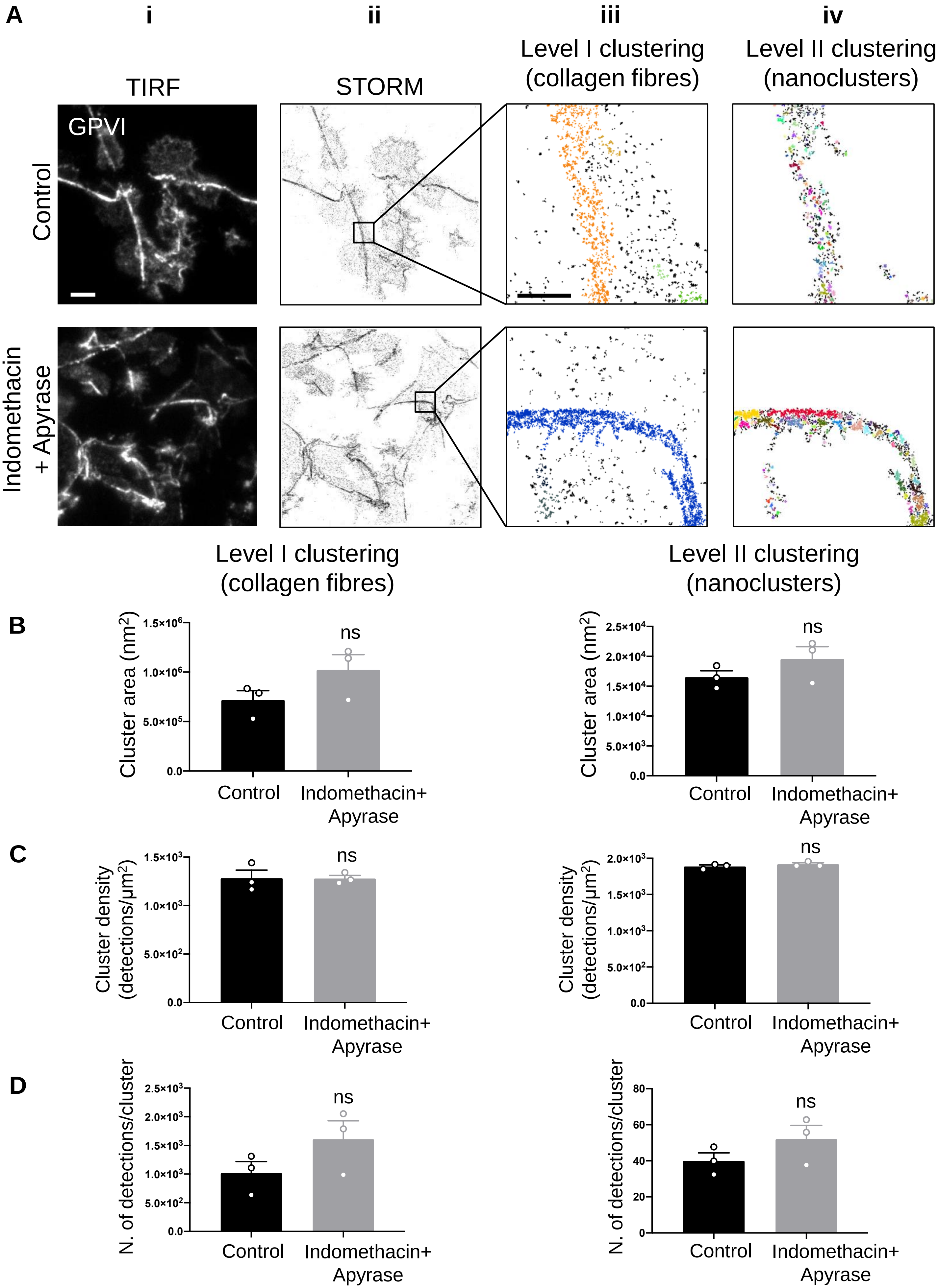


**Supp. Fig. 6 Inhibition of secondary mediators does not perturb GPVI cluster formation in platelets**

### Supplementary Fig. 7

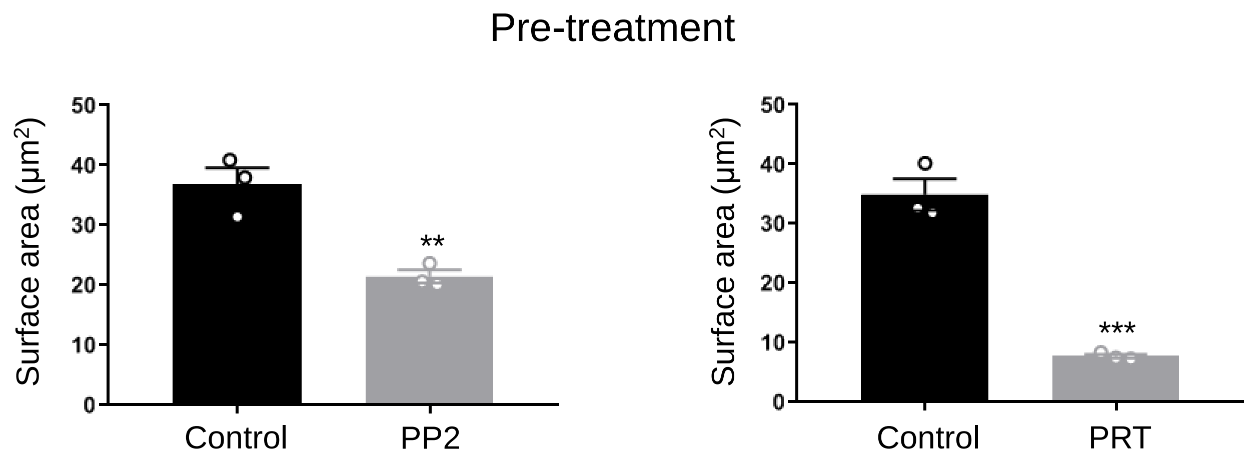


**Supp. Fig. 7 Effect of Src-family and Syk kinases of platelet spreading and GPVI clustering on collagen.**

Quantitative analysis of surface area in washed human platelets pre-incubated with DMSO (vehicle control), 20 µM PP2 (Src-family inhibitor) or 10 µM PRT (Syk inhibitor) and spread on collagen for 1 h. The graphs report the mean ± SEM from five FOVs from three independent experiments. The significance was measured using unpaired two-tailed t-test with ** $p < 0.01$ and *** $p < 0.001$.

#### Supplementary Fig. 8

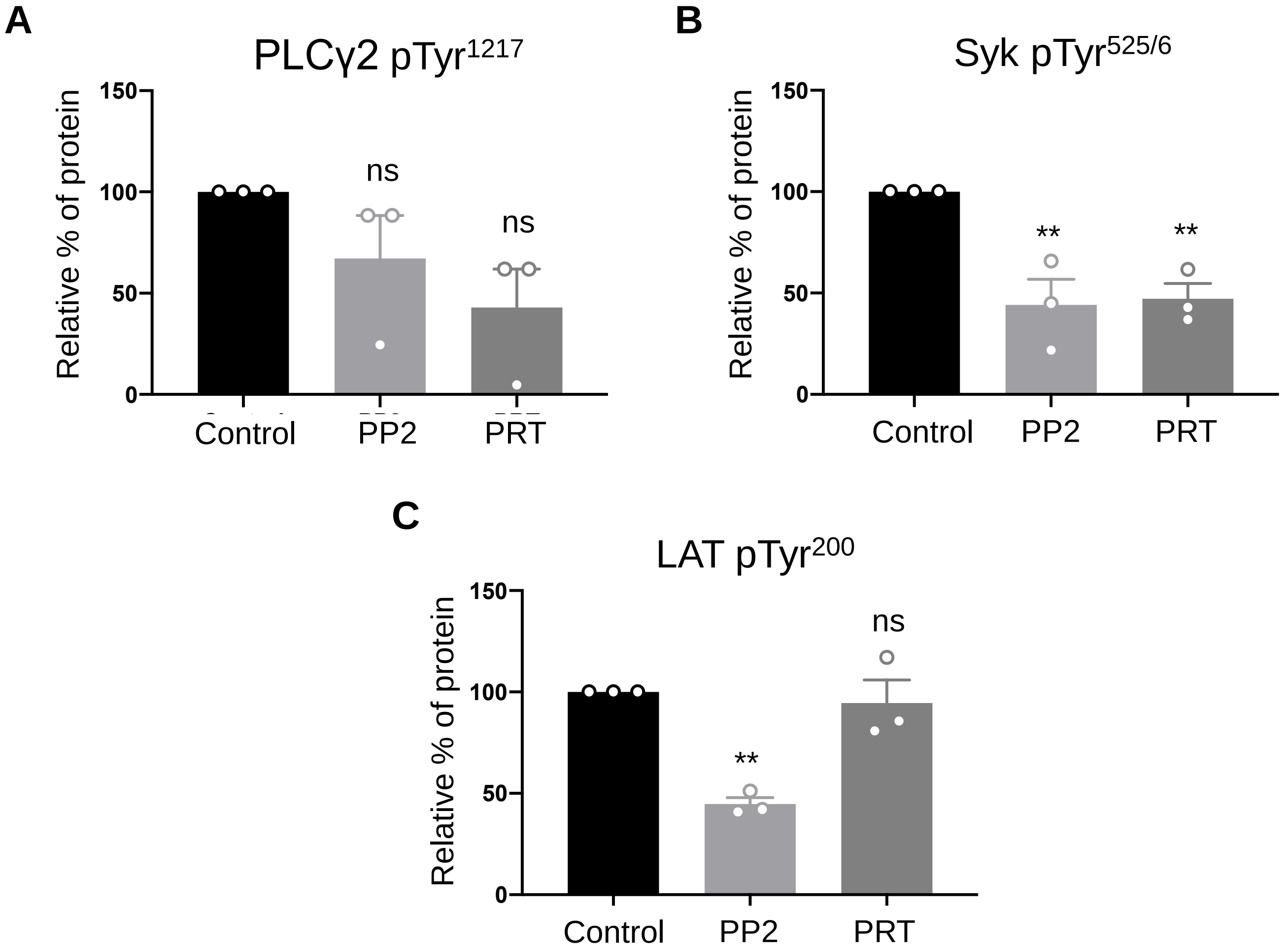


**Supp. Fig. 8 Western blot quantitative analysis of phosphorylated PLCγ2, Syk and LAT from lysates of platelets spread on collagen and and post-treated with Src-family and Syk inhibitors.** Quantification of band intensities of phospho-PLCγ2 (**A**), phospho-Syk (**B**) and phospho-LAT (**C**) normalised against the pan signals from lysates of platelets spread on collagen for 45 min and treated with DMSO (vehicle control), PP2 or PRT for 15 min (post-treatment). Data are expressed as mean ± SEM relative % of protein (Control was set at 100%) from three independent experiments. The significance was measured using one-way ANOVA with Tukey's multiple comparisons test with ** $p < 0.01$ and ns: not significantly different.

#### Supplementary Fig. 9

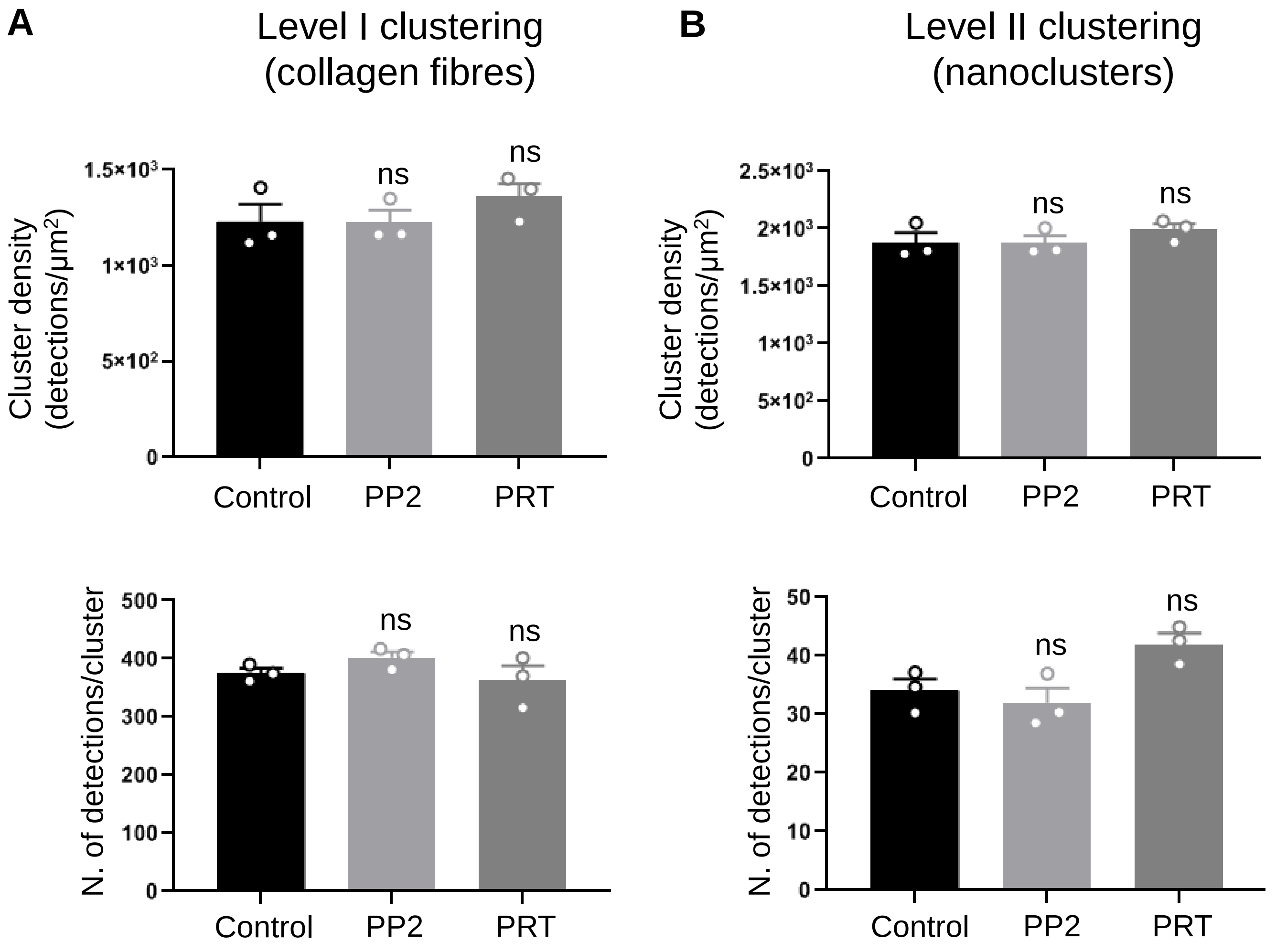


**Supp. Fig. 9 Two-level DBSCAN cluster analysis of GPVI in platelets spread on collagen and post-treated with Src-family and Syk inhibitors.**

Two-level DBSCAN cluster analysis of dSTORM images labelled for GPVI in platelets spread on collagen for 45 min and treated with DMSO (vehicle control), 20 μM PP2 or 10 μM PRT for 15 min (post-treatment). Quantification of GPVI cluster density and number of detections per cluster for both Level I (**A**) and Level II (**B**) clustering in control and treated platelets. Data are shown as mean ± SEM three independent experiments. One-way ANOVA with Tukey’s multiple comparisons test was used to calculate the statistical significance ($p < 0.05$; ns: not significantly different).

**Supplementary Fig. 10**

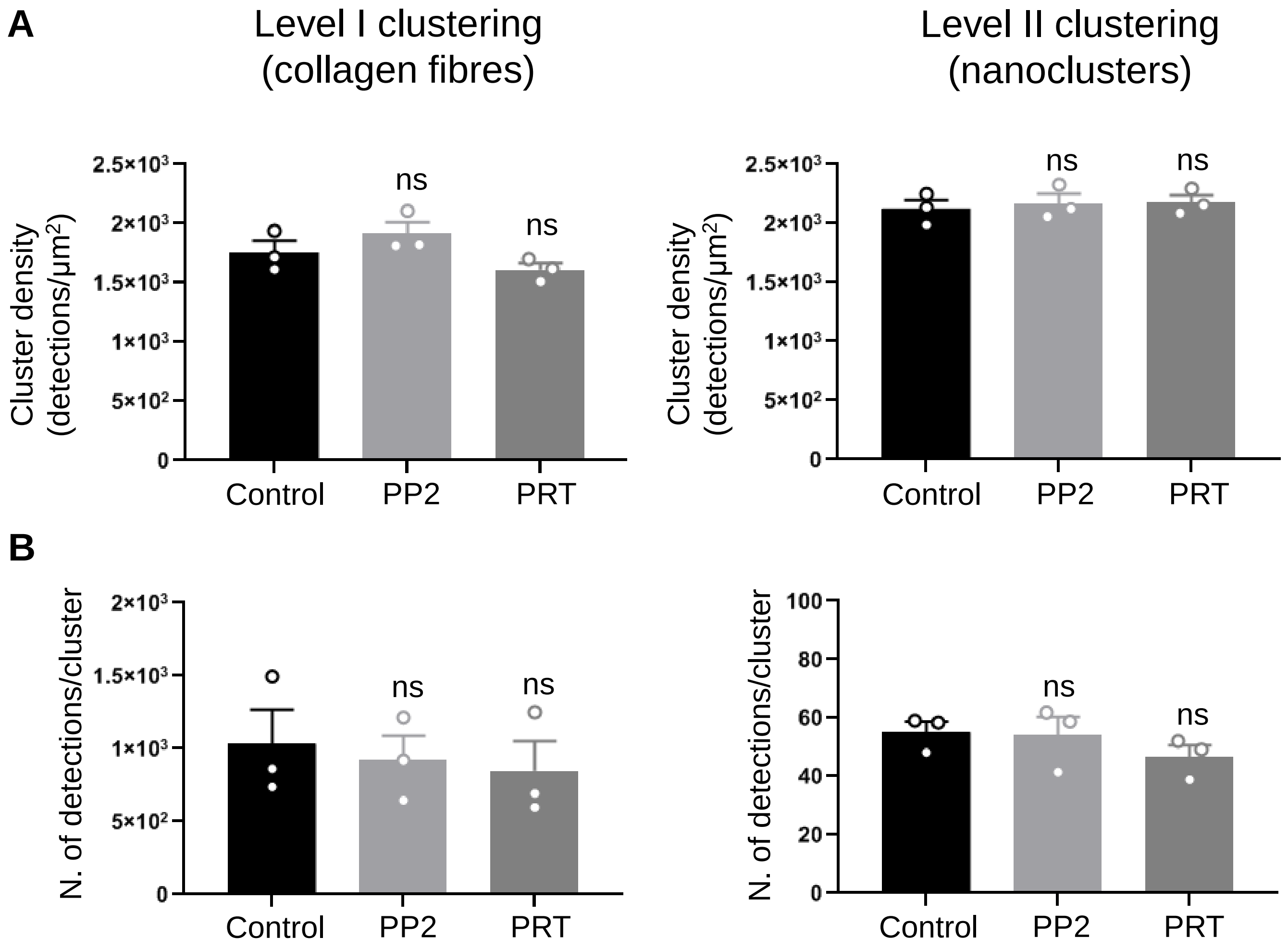


**Supp. Fig. 10 Two-level DBSCAN cluster analysis of integrin α2β1 in platelets spread on collagen and post-treated with Src-family and Syk inhibitors.**

Two-level DBSCAN cluster analysis of dSTORM images labelled for integrin α2β1 in platelets spread on collagen for 45 min and treated with DMSO (vehicle control), 20 µM PP2 or 10 µM PRT for 15 min (post-treatment). Quantification of integrin α2β1 cluster density (**A**) and number of detections per cluster (**B**) for both clustering levels in control and treated platelets. Bars represent mean ± SEM from three independent experiments. Significance was measured using one-way ANOVA with Tukey's multiple comparisons test ($p < 0.05$; ns: not significantly different).

### Supplementary Movies

**Supplementary Movie 1: Calcium mobilisation in platelets spread on collagen for different time periods.**
Live cell imaging of washed human platelets loaded with the $Ca^{2+}$ dye Oregon green 488 BAPTA-1-AM that had been spread on immobilised collagen for 15 min (top left panel), 30 min (top right panel), 1 h (bottom left panel) and 3 h (bottom right panel). Images were acquired every 1 sec for 2 min using a Zeiss Axio Observer 7 Epifluorescence microscope. Scale bar: 10 µm.

**Supplementary Movie 2: Calcium mobilisation in spread platelets after Src or Syk kinase inhibition.**
Live cell imaging of washed human platelets loaded with the $Ca^{2+}$ dye Oregon green 488 BAPTA-1-AM that had been spread on immobilised collagen for 45 min and then treated with either vehicle control, 20 µM PP2 (Src inhibitor) or 10 µM PRT (Syk inhibitor). Images were acquired every 1 sec for 2 min using a Zeiss Axio Observer 7 Epifluorescence microscope. Scale bar: 10 µm.
